# A Curvature Guided Composite Kernel Framework for Differential Gene Selection in Cancer Transcriptomics

**DOI:** 10.64898/2026.07.23.740443

**Authors:** Manan Gupta, Arka Prava Sarkar

## Abstract

Identification of differentially expressed genes is a crucial step for downstream tasks on gene data such as biomarker discovery, drug target identification. Traditional Methods assume negative binomial distribution on RNA-sequence data and models the DEGs using either generalized linear models or by estimating dispersion and assumption of mean-variance rate. The proposed method uses axiomatic approach by using quantum mechanics principles to project transcript data onto a Hilbert space using a composite kernel. Using the curvature generated by the transcripts on the latent manifold within the Hilbert space, a gravitational search inspired mechanism is used to identify the optimal number of differentially expressed genes by minimizing a representational loss function, and a reduced gene feature space is constructed as the potential differentially expressed genes. The proposed method has been compared with existing empirical methods for validation using proper statistical and biological benchmark analysis.

## Introduction

Differential expression (DE) analysis identifies genes whose expression levels change between two experimental conditions, typically disease and control/healthy conditions.^1^ Advancements in genome sequencing technologies, with the advent of next generation sequencing, have enabled genome-wide studies and analyses of differential expression.^2,3^ Differential expression analysis plays a pivotal role in the downstream process of biomarker prediction for complex diseases such as cancer.^4^ Differential expression analysis also helps identify driver and regulatory genes across biological pathways related to disease development and progression.^5^ These insights provide important insights into the disease and help in the development of personalised therapeutic solutions. There are well-established tools for differential expression analysis from both bulk NGS RNA-seq datasets and single-cell sparse transcriptomics datasets.^6^ Methods such as DESeq2 and edgeR rely on parametric models using the negative Binomial distribution, which assumes a count distribution. The existing tools handle the shrinkage estimator and false discovery rate control. The non-parametric approaches are mostly designed for sparse single-cell RNA sequencing (scRNA) datasets. SCDE and GLM-PCA explicitly model zero inflation and heterogeneity, but their drawback is handling sparsity in data, dropouts, and large scale data.

Robinson et. al. introduced the edgeR method, which models RNA-seq counts using a negative binomial distribution.^7^ They have used library-size normalization factors and estimated a common dispersion parameter across genes. After the initial estimation process, the edgeR method estimates gene-specific or tagwise dispersion, and an empirical Bayes shrinkage strategy is then used to pull the genes toward a common dispersion, thus improving stability when counts or replicates are limited at source. The generalised linear models used are mostly based on natural or quasi-likelihood frameworks, depending on design complexity. The limitations of edge are related to sample size and class imbalance. Strong heterocedasticity is unaccounted for by the dispersion model. Sometimes severe violations of the negative binomial mean-variance also affect downstream findings. Love et al. introduced the DESeq2 method, which uses shrinkage estimation to handle dispersion and fold changes, aiming to improve the stability and interpretability of the posterior estimates.^8^ Their method uses a generalised linear model to fit each gene iteratively. The read counts are then modelled using a negative binomial distribution with mean and dispersion parameters. Finally, a logarithmic link is used along with the GLMs where the method predicts the fold change score, which in turn is used to identify the DEGs. The limitations of the DESeq2 method lie in the domain of class imbalance and nonconformity to modeling sparse genetic data, such as scRNA-seq expression data. As with other common NB-based methods, sensitivity to outliers is a serious issue when predicting p-values for differential expression. Without prior robust settings, reduced power with extremely low replicate numbers also hinders DEG identification. Later in 2014, Law et al. and in 2015, Ritchie et al. introduced voom-limma to address issues with predecessor methods.^9,10^ The Limma method transforms RNA-seq counts to log-counts per million before estimating the mean-variance relationship across genes. The voom step converts the mean-variance trend; precision weights are calculated for each experimental observation, and the limma linear model is then used to identify the DEGs. Weighted linear models are fitted to each gene, and then an empirical Bayes shrinkage of residual variance is used to improve inference. It is also used to facilitate complex design variations, contrasts, and random-effect-style analytics. Voom-limma also provides similar detection performance to other methods. However, its limitation stems from assuming that the voom-derived weights can capture the mean-variance relationship adequately. This approach can be less robust if the sample size is small or the library or assay specification is highly heterogeneous. Apart from the discussed methods, another class of methods exists for the identification of DEGs from bulk RNA-seq count matrices. The second class is the non-parametric methods developed for DEG identification. Li and Tibshirani introduced SAMseq, a non-parametric method that utilised Wilcoxon-type test statistics integrated with resampling to handle variable sequencing depths.^11^ An estimator is used by the method, either by applying permutation or resampling, to obtain significant empirical null distributions and control false discovery rates without a prior assumption about a specific count distribution. SAMseq can be applied to both binary-class and multi-class RNA-sequence samples. The key limitation of SAMseq is the need for a moderate number of samples per experimental condition to achieve good sensitivity and low FDR. Also, heavy use of resampling makes SAMseq computationally expensive and not highly feasible for large datasets or complex covariate structures. Another notable non-parametric method, NOIseq, identifies DEGs by first building an empirical ‘Noise’ distribution of count changes by comparing samples from the same experimental condition (i.e., disease /normal). An assessment is then performed by the method to determine whether the observed changes between conditions exceed the noise distribution.^12^ The method includes variants for different experiments with and without biological replicates. This method is robust to sequencing depth variation and can control the FDR. However, it is limited in the raw output of the number of DEGs identified, resulting in significant data loss and missing biological insight during downstream analysis.

With the advancement of NGS technology, single cell sequencing has also become more efficient, enabling single-cell RNA-seq to scan and sequence RNA transcripts and yield high-throughput expression counts for thousands of cells collected from disease samples. scRNA-seq data, however, have one key difference from bulk RNA-seq data derived from multiple samples. scRNA-seq data are, by default, sparse. Therefore, parametric and non-parametric methods developed for bulk RNA-seq data do not perform well on scRNA-seq expression matrices.^13^ To overcome the limitations, several methods were developed that can handle not only the large volume of transcripts and cells but also the inherent sparsity of the scRNA-seq representation. Single cell differential expression (SCDE) uses a Bayesian framework to model scRNA-seq counts as a composite object comprising an expression component and a dropout component, each containing cell-specific error models.^14^ The method estimates posterior distributions for the mean expression level in each cell cluster and for fold changes, these parameters are used to compute the differential expression probabilities for each gene. By explicitly modelling dropouts and handling technical noise, SCDE shows higher sensitivity than bulk RNA-seq methods. The limitation of SCDE lies in its computationally intensive Bayesian modelling, and it often scales poorly with more modern, high-volume datasets. Further, Model based Analysis of single-cell transcriptomics, commonly known as MAST, has been conceptualised as a two-part hurdle model: logistic regression is used to estimate the probability of a gene’s expression, and a Gaussian linear model is used to model log-expression among detected cells.^15^ MAST enables integrative processing, incorporating covariates such as condition, batch, cellular detection rate, etc., into both components, which then jointly handle dropouts and continuous expression levels. The method is dependent upon hurdle model assumptions and treats cells as independent, which may induce pseudoreplication if the subject structure is ignored. Double hurdle modelling can become computationally expensive with increasing data volume; thus, scaling becomes a bottleneck when many biological replicates are available. The next widely used developmental method for modelling scRNA-seq data to identify DEGs is the NEBULA mixed-model method for multi-subject scRNA samples, which is capable of modelling both the subject-level and cell-level dispersions.^16^ A large-sample analytical approximation method is used for the marginal likelihood to avoid high-dimensional numerical integration, which enables better fitting to large droplet-based datasets. The NEBULA approximation relies on having many cells per subject; therefore, a performance drop is observed when cell counts are small or the design is imbalanced. The assumption of an appropriate negative binomial mean-variance structure for the data distribution and a random-effects specification can often lead to misspecification in some modern, high-feature-space datasets.

Kernel methods are used to embed data points into a Reproducing Kernel Hilbert Space (RKHS), allowing computational methods to capture nonlinear relationships present in the data.^17^ Formally, a kernel function is an inner product operation of the form K(x, y) = ϕ(x) · ϕ(y) that projects non-linearly separable data into a high-dimensional feature space. The most commonly known kernel function is the Linear kernel, which is the inner product k(x, x*^′^*) = x*^T^* · x*^′^* that corresponds to the identity feature map into a finite dimensional Hilbert space.^18^ The linear kernel induces an RKHS considered with the original Euclidean space. The limitations of this method include the inability to retain the vast biological knowledge encoded in non-linear, many-dimensional embeddings latent in high-volume, high throughput next generation sequencing (NGS) data. The polynomial kernel corresponds to an explicitly mapped finite-dimensional space of monomials up to degree d, which forms an RKHS of polynomial functions.^19^ Though the degree can be selected explicitly, the RKHS formed still remains finite dimensional, thus becoming less suitable for representing the rich, interconnected, non-linear, non-hierarchical relationships among multiple genetic data points available in NGS data. The Gaussian kernel or the radial basis dunction (RBF) can define an infinite dimensional RKHS of very smooth functions and a strictly positive-definite kernel.^20^ The smooth functional RKHS, despite being very efficient at modelling time course changes or spatial trends in biological data, often loses sharp biological features due to transcriptional bursts, threshold effects, and oversmoothing. Using too few basis functions results in biased modelling of data, while using too many causes the model to hallucinate. Boundary effects can also become prominent when using RBF kernels, where smoothers become highly unstable at the separation line between trends present in genetic datasets.

The proposed method introduces a new paradigm for DEG identification based on a quantum mechanical analogy. The method is designed as a modality-agnostic process that can perform DEG selection without depending on the data type, whether the data falls under the bulk RNA-seq modality or scRNA. The proposed methodology uses a composite kernel to project transcriptomic expression matrix into a Hilbert space. It uses a gravitational search inspired algorithm on the curvature of transcripts produced on the embedding manifold to determine and select the optimal number of differentially expressed genes. The representational loss function ensures that significant data loss has not occurred while performing the selection. The results have been experimentally validated by comparing with existing methods for DEG selection and shows satisfactory results both in terms of classification performance and biological enrichment analysis of the identified genes.

## Materials and Methods

The standard time independent Schrödinger equation is given by **H** |Ψ⟩ = E |Ψ⟩.^21,22^ In this work, we describe the gene’s state using the first postulate of quantum mechanics, i.e., it is a physical state of system in Hilbert space described by the wave function |Ψ⟩*_gene_*. Using the inner product rule of wave functions in Hilbert space, the following condition is imposed on the gene state wave function

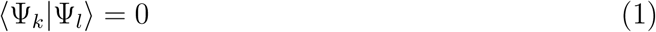

which implies that the information of two different genes are orthogonal to each other, i.e., no information of one gene state is present in the other gene state. Using the method of linear combination of atomic orbitals (LCAO), the gene state wavefunction can be written in the form of linearly independent basis sets of the following form:

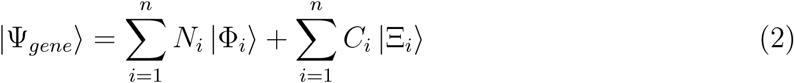

Equation 2 establishes the Composite Nature of the framework. Unlike standard feature vectors where dimensions are treated uniformly, this state vector is structurally partitioned into two distinct regimes: the normal regime spanned by the wavefunctions |Φ*_i_*⟩, and the cancer regime spanned by the wavefunctions |Ξ*_i_*⟩. The coefficients N*_i_* and C*_i_* represents the weight factor magnitudes the gene expression in the normal and cancer contexts, respectively. In this work, we are considering 50 different genes, for which 50 different normal cell regime, and 50 different cancer cell regimes are considered for each gene wavefunction. This also implies that a gene is not a single point in space but a dual entity capable of simultaneous projection into healthy and pathological wavefunctions. The Composite Kernel must, therefore, be capable of ingesting data that preserves this duality.

In order to ensure that equation 1 is consistent for all the genes, the following axioms are imposed on the normal and cancer cell wavefunctions:

Axiom I: The normal cell and the cancer cell wavefunctions are always orthogonal to each other. This is a direct consequence that arises from the LCAO condition, as we represent the gene wavefunctions in terms of normal and cancer cell wavefunctions. This implies that these wavefunctions are linearly independent as well. Mathematically, ⟨Φ|Ξ⟩ = 0

Axiom II: The basis sets spanning the normal cell and cancer cell wavefunctions are orthogonal to each other. This can be represented mathematically as ⟨Φ*_i_*|Φ*_j_*⟩ = 0 and ⟨Ξ*_i_*|Ξ*_j_*⟩ = 0.

Let an expression matrix be

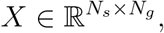

where N*_s_* is the number of samples and N*_g_* is the number of genes. Labels

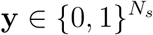

partition the samples into normal (0) and cancer (1) regimes. For gene g, define the coefficient vector

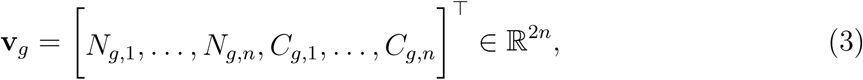

where N*_g,i_* and C*_g,i_* are the expression magnitudes of gene g in the normal and cancer regimes, respectively. Here, n denotes the number of replicates or pooled samples per regime in the manuscript notation; in the implementation, n indexes the samples within each class. The input transformation matrix is then obtained using the condition T (v) = **P** = v*^T^* v. The resulting matrix **P** is a square matrix of dimensions 2n × 2n where P*_ij_* = v*_i_*v*_j_*.

For gene g, with coefficient vector **v***_g_*, define the composite kernel functional as

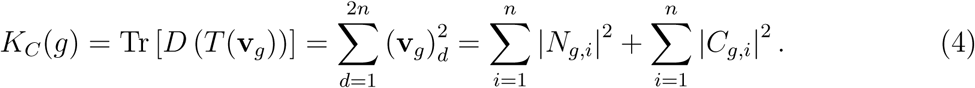

For implementation on a full matrix X, for each column (gene) g:

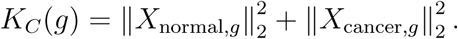

**Definition 1** (Composite inner product). *For* **u**, **w** ∈ R^2n^, *define*

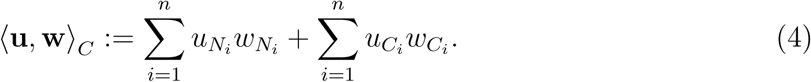

**Lemma 1.** ⟨·, ·⟩*_C_ is a valid inner product on* R^2n^.

*Proof.* Bilinearity and symmetry follow from the definition. Positive definiteness:

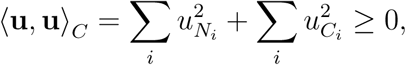

with equality iff all components vanish. Hence it induces a Hilbert space H*_C_*.

**Theorem 1** (PSD of the composite kernel Gram matrix). *Let*

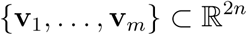

*be coefficient vectors of* m *genes. Define the Gram matrix* G ∈ R*^m^*^×*m*^ *by*

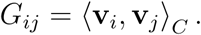

*Then* G *is symmetric and positive semi-definite*.

*Proof.* Symmetry is immediate. For any **a** ∈ R*^m^*,

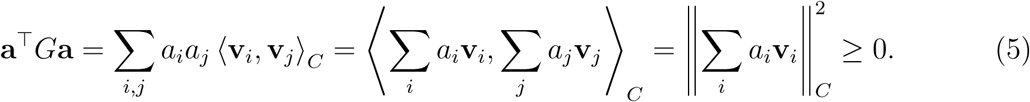

Thus G ⪰ 0. The scalar gene score (3) satisfies

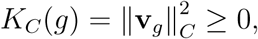

consistent with the diagonal of the PSD Gram matrix when genes are orthonormal in feature space.

The Kernel obtained from equation 4 fundamentally differs from RKHS and other machine learning based kernels in the fact that a composite kernel in standard machine learning algorithms is the linear combination of multiple kernels (K = *_m_* µ*_m_*K*_m_*). The framework presented in this paper represents the state vector itself, which is not the typical scalar addition of functions.

The following is then the algorithm implemented to implement the Kernel for a particular same of gene, say k:

- Construct v*_k_* ∈ R^100^
- _•_ Compute **M***_k_* = v*^T^* v*_k_*
- Set M*_ij_* = 0 for i ̸= j by applying the mask operator
- Compute the sum

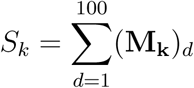

A positive-definite kernel naturally gives rise to a reproducing kernel Hilbert space through the Aronszajn construction.^23^ When the underlying domain has the structure of a smooth manifold, the kernel-induced feature map places this manifold inside the RKHS. The inner product of the RKHS can then be pulled back to the original manifold, giving it a Riemannian metric.^24–26^ Therefore, the Hilbert space generated by the kernel from equation 4 can be modeled as a Riemannian manifold M as well. The intrinsic geometry of this manifold, including concepts of distance, angles, and curvature, is entirely determined by its metric tensor g*_ij_*. For a strictly diagonal metric tensor where g*_ij_* = 0 when i ̸= j and where the contravariant metric tensor is simply the inverse of the diagonal elements g*^ii^* = (g*_ii_*)^−1^, the Christoffel symbols reduce to a highly simplified set of non-zero components.^27^ The Riemann curvature tensor R_σμν_*^ρ^* which encodes all the information regarding the intrinsic curvature of the manifold, is computed from the spatial derivatives of the Christoffel symbols.^28^ For an arbitrary diagonal metric, the diagonal components of the Ricci tensor can be efficiently computed without calculating the full Riemann tensor, utilizing formulations that depend solely on the logarithmic derivatives of the metric components. ^28^ This mathematical simplification guarantees that the geometric analysis remains computationally efficient and mathematically exact, allowing for the scaling of the algorithm to genome-wide datasets without succumbing to exponential time complexity. Although the continuous formulation of the Ricci tensor provides the theoretical underpinning for the manifold’s geometry, real-world gene expression data consists of discrete point clouds sampled from the underlying biological distributions. ^29^ Applying continuous differential operators directly to noisy, finite datasets is computationally unstable. Therefore, an estimator for discrete Ricci curvature is required to approximate the continuous geometry.^30^ In this work, we implement the Oliver-Ricci curvature (ORC) as it relies heavily on the discrete topology of the network rather than continuous geometric distances.^31^ The mathematical form of the ORC is given by the equation 5 computed over the local neighborhood N(x) of the gene, weighted by the transition probabilities

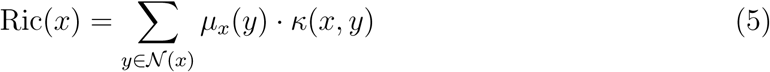

where µ*_x_*is the probability measure defining a random walk centered at gene x, and κ(x, y) is the Oliver-Ricci curvature.^31,32^ This aggregation process collapses the local geometric structure into a single density value for each gene, mapping the complex topological dynamics into an interpretable spectrum.

While the topological visualization of the Ricci curvature density successfully isolates the geometric anomalies associated with DEGs, pure visualization is intrinsically subjective. Relying on visual clustering or manual thresholding to determine the exact cutoff point for feature selection is suboptimal, as it fails to quantify the precise trade-off between dimensionality reduction and information retention.^33^ To autonomously extract the optimal number of DEGs (k) without relying on visual heuristics, a systematic mathematical method to quantify and minimize data loss is required. To fulfill this requirement, a novel, dynamic physics-inspired optimization algorithm is developed. Drawing upon the mechanics of the gravitational search algorithm (GSA) proposed by Rashidi et al. and recent advancements in physics-informed neural networks (PINNs) used for gravity field modeling.^34,35^ This framework simulates a dynamic gravitational potential field within the Hilbert space. This physical simulation dynamically orchestrates the feature space to minimize a mathematically rigorous representational data loss. The mathematical formulation of the data loss is given in the form

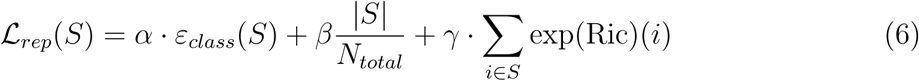

where ε*_class_*(S) represents the empirical classification error, 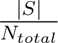 is a linear penalty for the inclusion of excessive features, promoting network sparsity and preventing overfitting, Σ*_i_*_∈_*_S_* exp(Ric)(i) is a geometric regularization term. Critical DEGs can exhibit highly negative Ricci curvature which leads to the exponential term to approach zero. Therefore, the loss function is heavily penalized for including flat or positively curved redundant genes, while the inclusion of negatively curved bottleneck genes incurs almost zero geometric cost. ^36–39^ α, β, γ are tunable scalar hyperparameters that control the relative weighting of classification accuracy, subset sparsity, and geometric anomaly inclusion, S is a candidate subset of genes with cardinality k = |S| and N*_total_* is the total number of genes in the dataset. The loss function defined in equation 6 is then implemented the gravitational search algorithm, where the dynamical mass update equation for a particular gene i is calculated iteratively by evaluating its individual fitness contribution toward minimizing the global representational loss using equation 7, which is similar to the formulation given by Su et. al.^40^

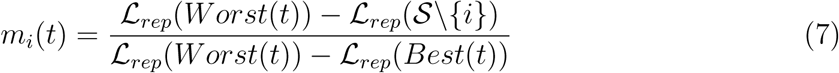

where L*_rep_*(S\{i}) evaluates the detrimental impact of removing gene i from the current subset. Genes whose removal causes a significant spike in loss (due to dropping classification accuracy or losing a critical geometric bottleneck) will yield a higher fitness score m*_i_*(t). The normalized active gravitational mass for the simulation is then computed using equation 8.

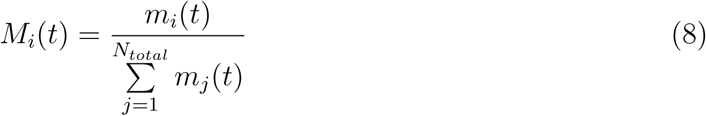

Finally, the gravitational force vector F*_ij_*(t) on gene i due to gene j is defined by their respective masses and the intrinsic geometric distance separating them which is shown in equation 9.^41^

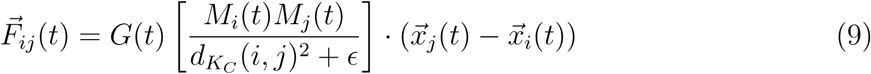

where d*_KC_* is the exact geodesic distance in the Hilbert space generated by the kernel, ɛ is a small stabilisation constant to prevent division by zero during particle collisions, x⃗ represents the n-dimensional spatial coordinates of the gene, G(t) is the dynamic gravitational constant.^40,41^ The complete workflow is shown in the Figure 1.

**Figure 1:**
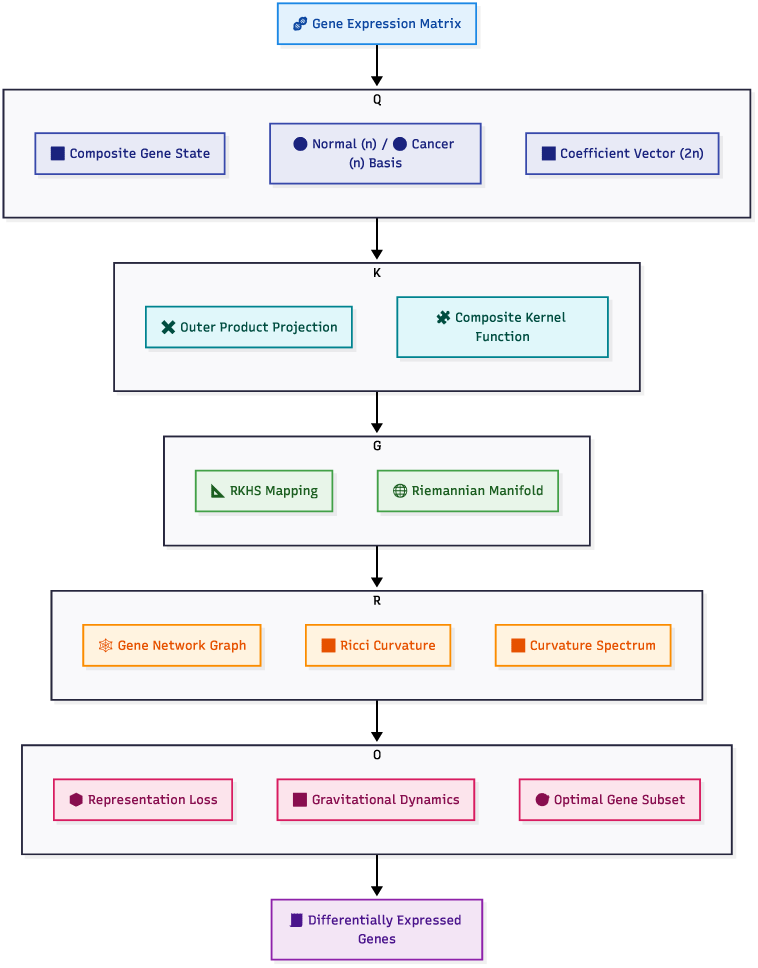
Complete workflow of the method presented.

The new method was rigorously tested using a standard classification experiment and biological enrichment analysis. This section will discuss the experimental pipeline, then present the statistical validation results. For experimental validation three TCGA NGS RNA sequence datasets and thee gene expression omnibus (GEO) datasets were used. ^42,43^ The dataset description is given in table 1. For the benchmarking, first every original raw datasets were cleaned by eliminating missing values and normalized using Z-score method. The preprocessed original datasets were then classified using support vector machine classification algorithm with radial basis function kernel, and XGBoost algorithm with 20 estimators. Each classification algorithm was iterated 50 times, the average value of the accuracy, precision, recall and F1 metrics were taken. Then from the original datasets all transcripts/genes were eliminated except the ones selected by proposed method. The support vector machine and XGBoost classifier was used to classify the reduced datasets using the same parameters used for the original datasets. The classification metrics empirically show that the selection of differentially expressed genes by proposed method is capable of delivering better results than using the original datasets except some minor exceptions. Table 1–4 below contains the descriptions of the datasets used to test the proposed method for both internal and external validation respectively.

**Table 1:**
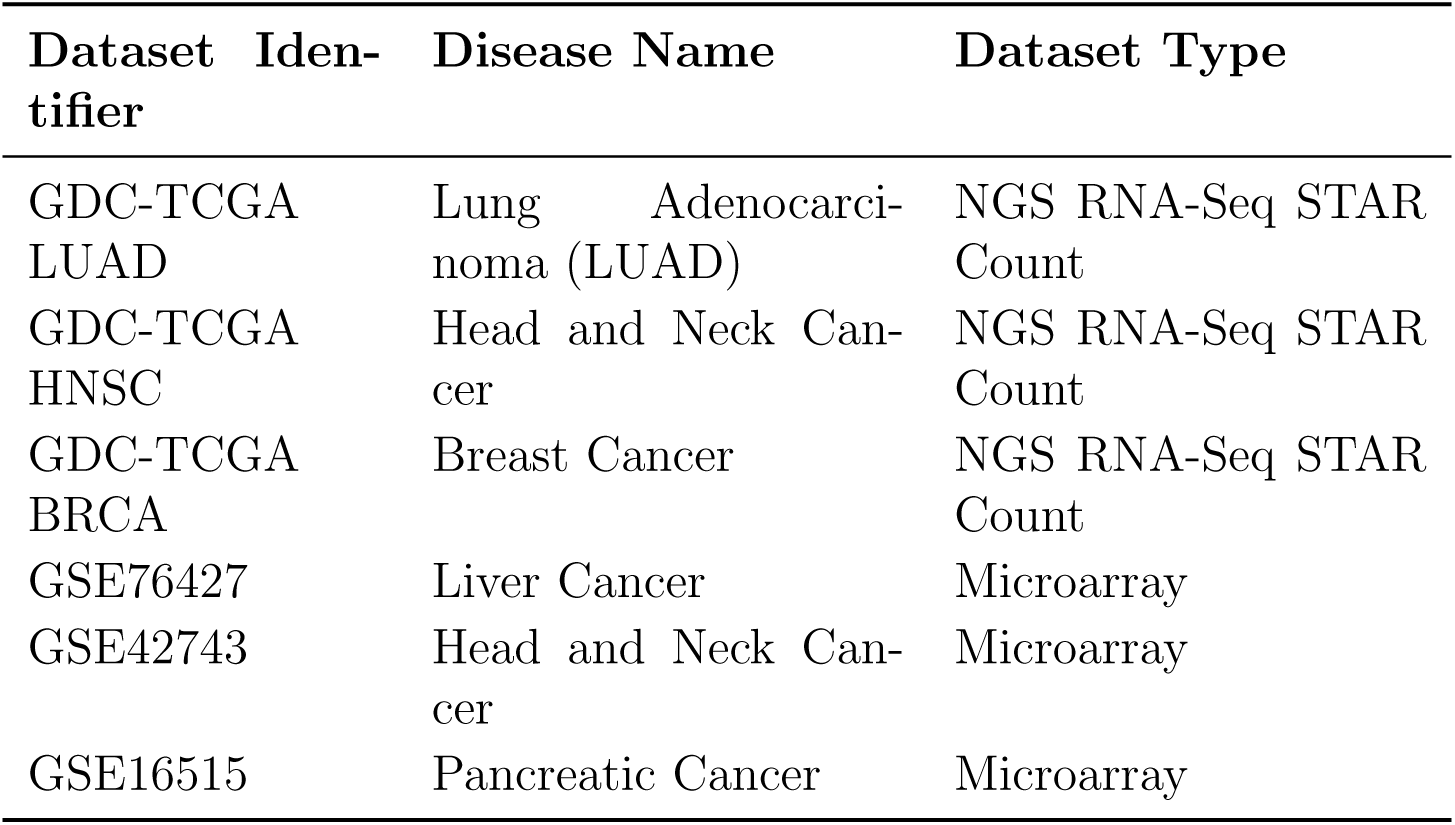
Summary of cancer gene expression datasets used in this study.

| Dataset Identifier | Disease Name | Dataset Type |
| --- | --- | --- |
| GDC-TCGA LUAD | Lung Adenocarcinoma (LUAD) | NGS RNA-Seq STAR Count |
| GDC-TCGA HNSC | Head and Neck Cancer | NGS RNA-Seq STAR Count |
| GDC-TCGA BRCA | Breast Cancer | NGS RNA-Seq STAR Count |
| GSE76427 | Liver Cancer | Microarray |
| GSE42743 | Head and Neck Cancer | Microarray |
| GSE16515 | Pancreatic Cancer | Microarray |

**Table 2:**
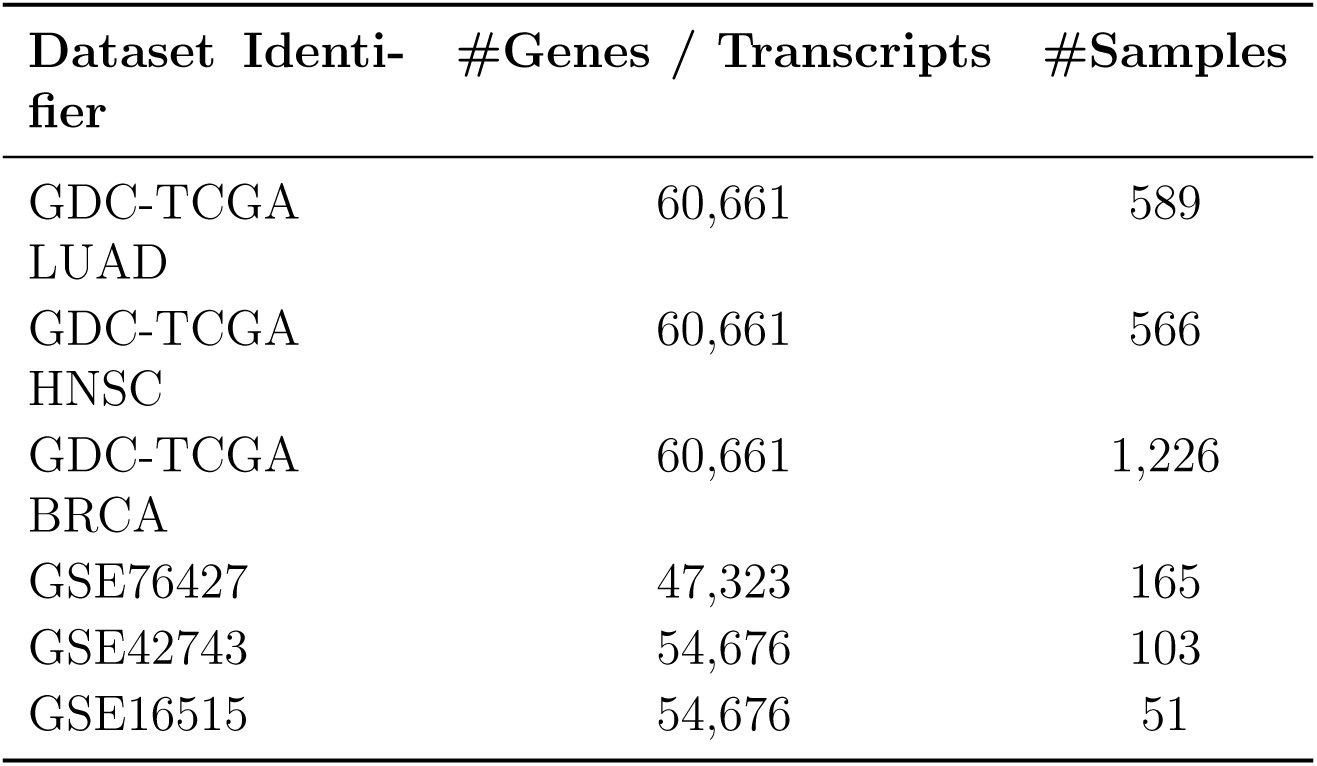
Gene/transcript and sample counts of the cancer gene expression datasets used in this study.

**Table 3:** Additional cancer datasets used for external validation and comparative analysis.

| Dataset Identifier | Disease Name | Dataset Type |
| --- | --- | --- |
| GSE19804 | Lung Adenocarcinoma | Microarray |
| GSE70947 | Breast Cancer | Microarray |
| GDC-TCGA LIHC | Liver Cancer | NGS RNA-Seq STAR Count |
| GDC-TCGA PAAD | Pancreatic Cancer | NGS RNA-Seq STAR Count |

**Table 4:**
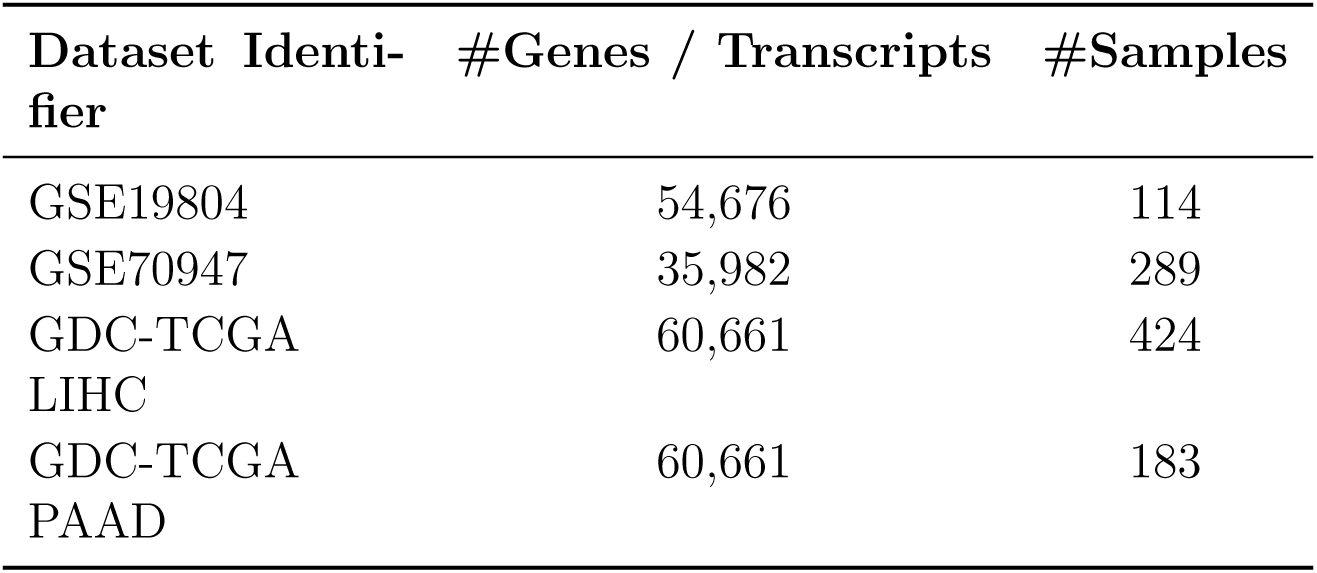
Gene/transcript and sample counts of the additional cancer datasets used for external validation and comparative analysis.

| Dataset Identifier | #Genes / Transcripts | #Samples |
| --- | --- | --- |
| GSE19804 | 54,676 | 114 |
| GSE70947 | 35,982 | 289 |
| GDC-TCGA LIHC | 60,661 | 424 |
| GDC-TCGA PAAD | 60,661 | 183 |

### Comparative Analysis

The same datasets were used to perform comparative analysis of existing differentially expressed gene identification methods. Two parametric methods DESeq2, edgeR and one non-parametric method NOISeq was used for the study. Each original dataset was processed by the three methods to identify the exact number of differentially expressed genes identified by the gravity function introduced by the proposed method. Once the differentially expressed genes were identified by the three chosen existing methods, three reduced forms of each dataset was constructed by the identified DEGs determined by each method, for every datasets used for the original statistical validation. Using the same classification experimental pipeline used for the original datasets, the reduced versions of the datasets were classified and accuracy and F1 score was used to compare the performance of the proposed method with DESeq, edgeR and NOISeq. The results show potential supereor performance in DEG selection by the proposed method as can be seen in table 5.

**Table 5:**
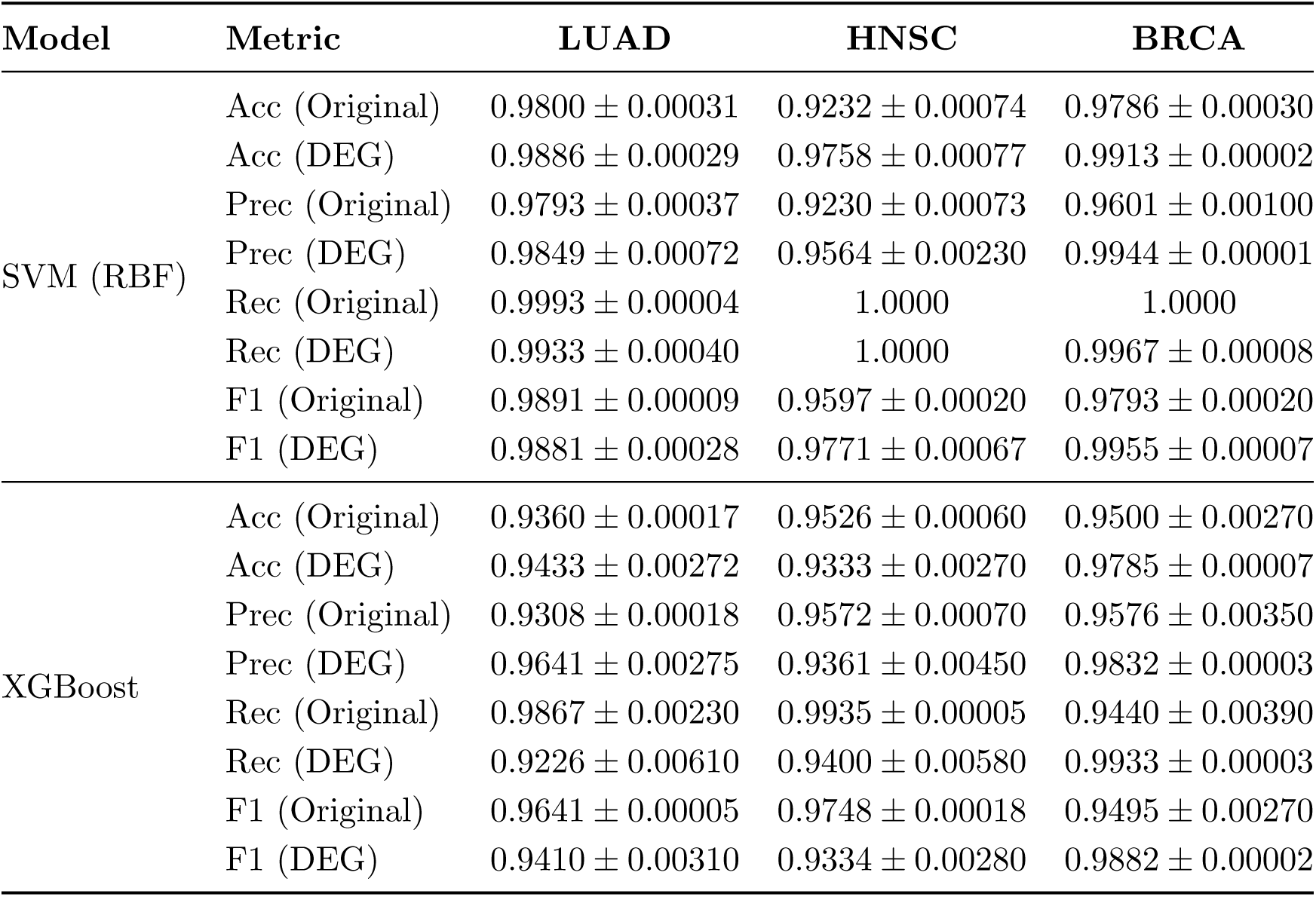
Performance comparison of SVM with RBF kernel and XGBoost models on LUAD, HNSC, and BRCA datasets using original and DEG-based feature sets.

## Results and Discussion

Table 5 shows the classification performance of the selected DEGs by the proposed method compared to the classification performance of the original unfiltered TCGA datasets. Table 6 shows the classification performance of the selected DEGs by the proposed method on CuMiDa datasets compared to the original microarray datasets. The observation clearly demonstrates that across the TCGA benchmark cohorts, the feature space of the selected DEGs has consistently improved discrimination relative to the original transcriptome for both the SVM and XGBoost classifier. The strongest gains were observed for SVM with RBF kernel. In all three TCGA cohorts LUAD, HNSC and BRCA classification accuracy increased measured by SVM was increased from 98%, 92.32% and 97.86% to 98.86%, 97.5% and 99.13% respectively, accompanied by higher precision, recall and F1 scores as shown in table 5. The results indicate that the proposed method selected genes that retained more tumor specific statistically relevant signals while reducing redundancy. XGBoost displayed a similar but modest improvement slope on TCGA cohorts, while exhibiting the most superior performance for BRCA where accuracy rose from 95% to 97.85% and F1 score from 94.95% to 98.82% as shown in table 5. The results show that the proposed method can extract statistically relevant DEGs from NGS RNA sequence datasets, while not reducing the quality of the transcriptomic feature space.

**Table 6:** Performance comparison of SVM with RBF kernel and XGBoost models on CuMiDa liver, throat, and pancreatic cancer datasets using original and DEG-based feature sets.

| Model | Metric | CuMiDa |  |  |
| --- | --- | --- | --- | --- |
|  |  | Liver | Throat | Pancreatic |
| SVM (RBF) | Acc (Original) | $0.9508 \pm 0.0006$ | $0.7690 \pm 0.0121$ | $0.70375 \pm 0.0085$ |
| | Acc (DEG) | $0.9455 \pm 0.0009$ | $0.7971 \pm 0.0118$ | $0.7055 \pm 0.0001$ |
| | Prec (Original) | $0.9543 \pm 0.0006$ | $0.6721 \pm 0.0488$ | $0.5167 \pm 0.2520$ |
| | Prec (DEG) | $0.9170 \pm 0.0025$ | $0.7343 \pm 0.4030$ | $0.4978 \pm 0.0002$ |
| | Rec (Original) | $0.9508 \pm 0.0006$ | $0.7688 \pm 0.0120$ | $0.7035 \pm 0.0860$ |
| | Rec (DEG) | $0.9455 \pm 0.0009$ | $0.7974 \pm 0.0121$ | $0.7061 \pm 0.0021$ |
| | F1 (Original) | $0.9512 \pm 0.0006$ | $0.6967 \pm 0.0280$ | $0.5881 \pm 0.1630$ |
| | F1 (DEG) | $0.9025 \pm 0.0011$ | $0.7277 \pm 0.0236$ | $0.5837 \pm 0.0002$ |
|  | AUC (Original) | – | – | – |
|  | AUC (DEG) | – | – | – |
| XGBoost | Acc (Original) | $0.8132 \pm 0.0245$ | $0.7271 \pm 0.0064$ | $0.6962 \pm 0.0067$ |
| | Acc (DEG) | $0.8933 \pm 0.0021$ | $0.8357 \pm 0.0042$ | $0.8455 \pm 0.0170$ |
| | Prec (Original) | $0.7230 \pm 0.7270$ | $0.5541 \pm 0.0213$ | $0.5001 \pm 0.0177$ |
| | Prec (DEG) | $0.8923 \pm 0.0042$ | $0.8395 \pm 0.0046$ | $0.8354 \pm 0.1980$ |
| | Rec (Original) | $0.8131 \pm 0.2450$ | $0.7271 \pm 0.0062$ | $0.6962 \pm 0.0067$ |
| | Rec (DEG) | $0.9013 \pm 0.0048$ | $0.8353 \pm 0.0049$ | $0.8451 \pm 0.0172$ |
| | F1 (Original) | $0.7555 \pm 0.0505$ | $0.6242 \pm 0.0138$ | $0.5780 \pm 0.0126$ |
| | F1 (DEG) | $0.8940 \pm 0.0022$ | $0.8307 \pm 0.0050$ | $0.3760 \pm 0.0192$ |
|  | AUC (Original) | – | – | – |
|  | AUC (DEG) | – | – | – |

Figure 2 shows the mean AUC and AUROC plots over 50 runs of both classifiers for all three TCGA cohorts. It clearly demonstrates that the DEG selected feature set consistently achieved high quality class discrimination along with no visible signs of overfitting or underfitting. The higher AUC values across all three TCGA cohorts indicate enhanced sensitivity-specificity balance and reduced feature redundancy.

**Figure 2:**
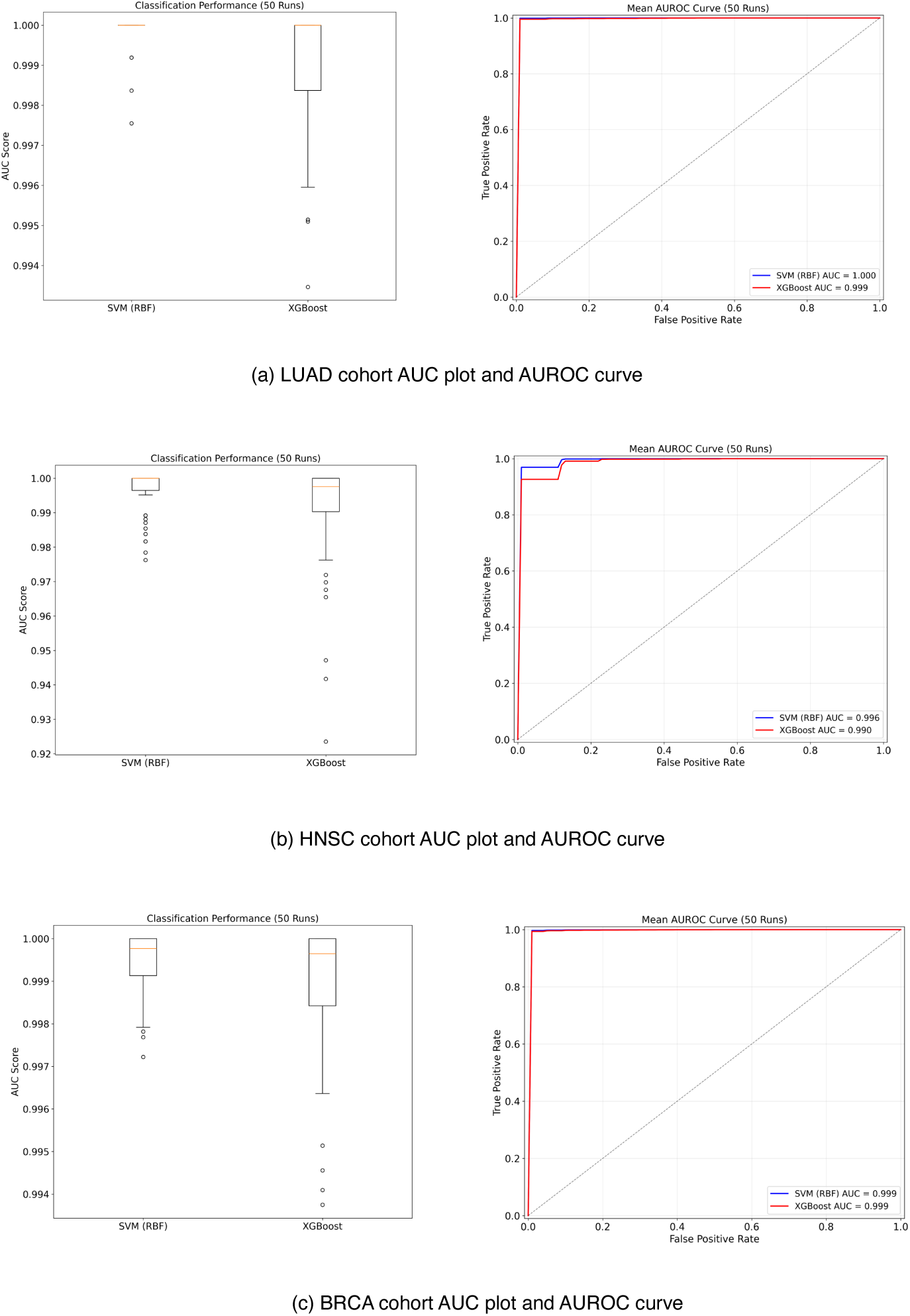
Mean AUC and AUROC plots of all three TCGA cohorts for 50 runs.

The same method was used to select DEGs from CuMiDa microarray datasets to evaluate modality agnostic nature of the proposed method. Though the effect of selected DEGs were more classifier dependent when the experimental pipeline was executed, it shows, that selecting DEGs by the proposed method in most cases retains the statistical significance compared to the original microarray datasets. Except for liver cancer dataset, which significantly dropped in classification accuracy by SVM. But using an ensemble classifier demonstrated that consistent performance gain was achieved throughout all three different tissue samples as shown in table 6. The results suggests that the proposed DEG selection method generalizes well across sequencing and microarray platforms.

Similarly as can be seen on the AUC and AUROC plots for the CuMiDa microarray cohorts, the DEG feature space improves classification robustness, specially for XGBoost method. Enhanced ROC characteristics and increased trends of AUC in the liver, throat and pancreatic cohorts also indicates improved discrimination capability and reduced interference from noisy microarray features. The plots are shown in Figure 3. These findings suggests that the proposed method is capable enough to generalize over microarray platform generated data as well.

**Figure 3:**
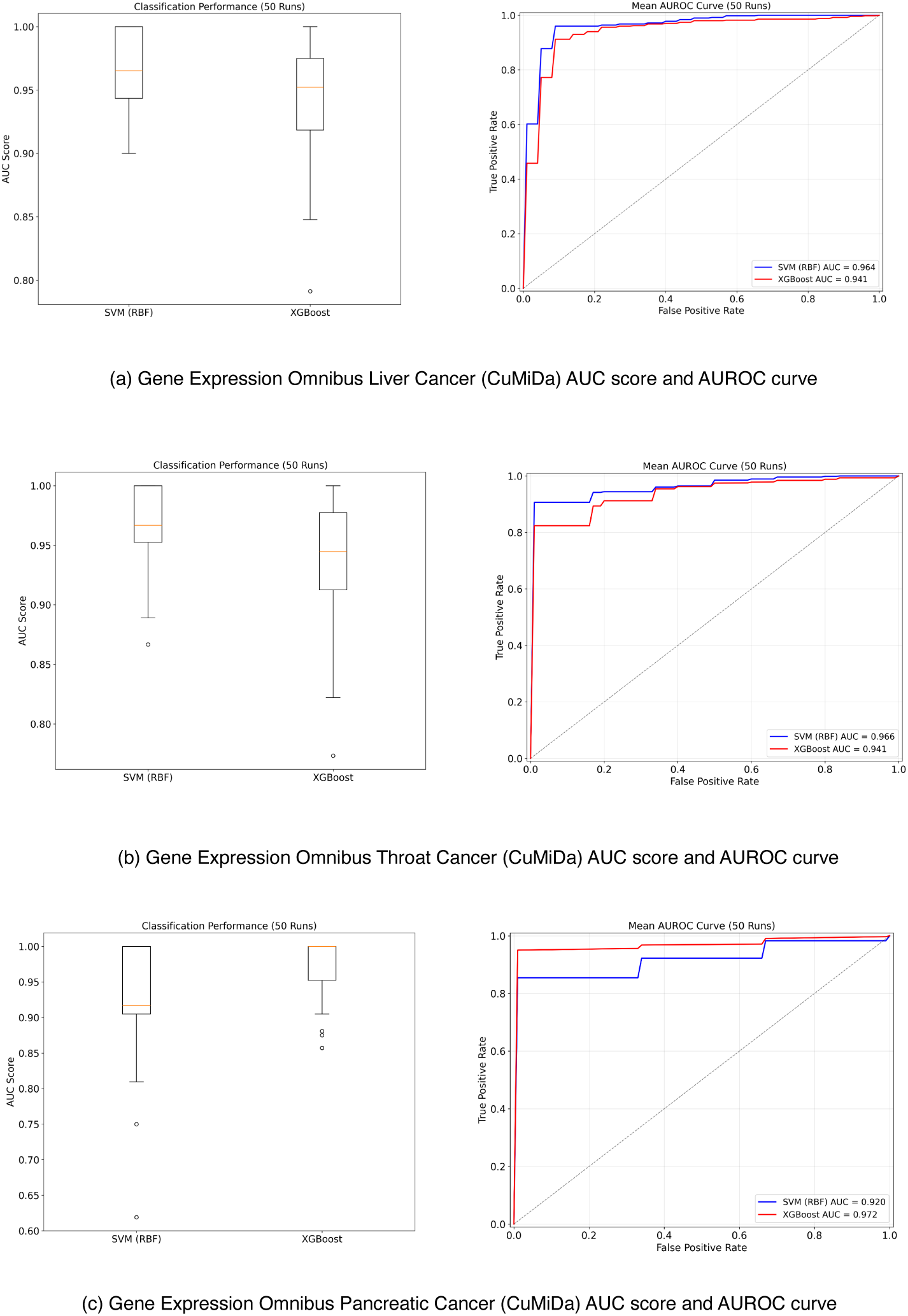
Mean AUC and AUROC plots of CuMiDa cohorts for 50 runs.

The methods capabilities were also analysed via latent space visualization and distribution analysis. The following figures 4 and 5 demonstrates and describes PCA and UMAP based component distribution analysis for the predicted DEGs for both the TCGA and CuMiDa cohorts respectively. It can be established from the PCA and UMAp embeddings of the TCGA cohorts that the proposed DEG selection framework generates highly discriminative low-dimensional feature manifold with improved intra-class compactness while exhibiting inter-class separability between gene expressions level-wise. The DEG derived representations have reduced cluster overlap and exhibits more coherent spatial organization in all three TCGA cohorts. These findings indicates the effective elimination of redundant and noisy transcriptomic features while preserving informative variance. Additionally, the concordance between the linear PCA and non-linear UMAP manifolds suggest that the selected DEGs retain both global variance structure and local neighbourhood topology, this validates the robustness and stability of the extracted feature space for downstream cancer subtype classification and biomarker discovery.

**Figure 4:**
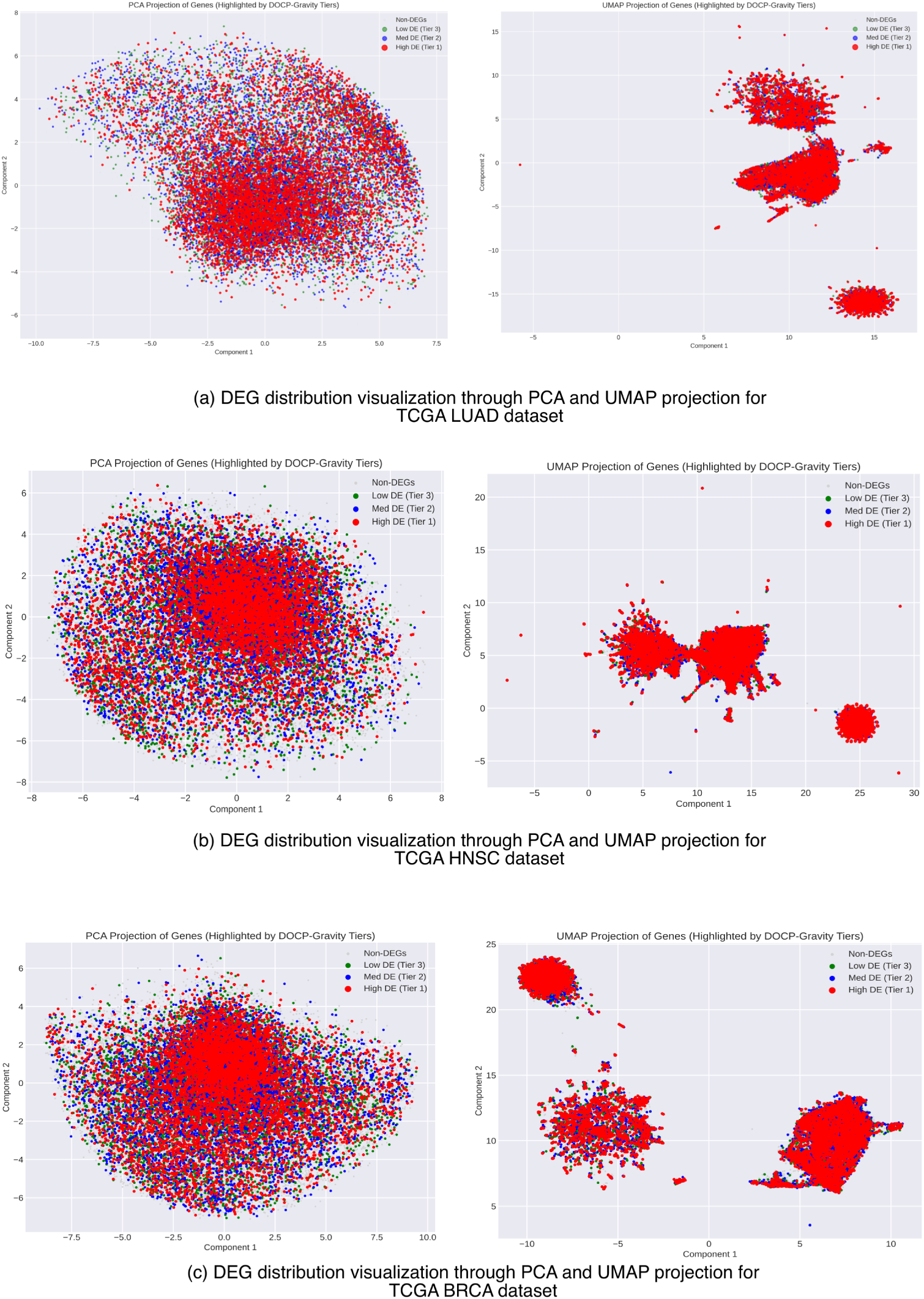
Visualisation of DEG distribution and UMAP projection of LUAD, HNSC and BRCA datasets.

**Figure 5:**
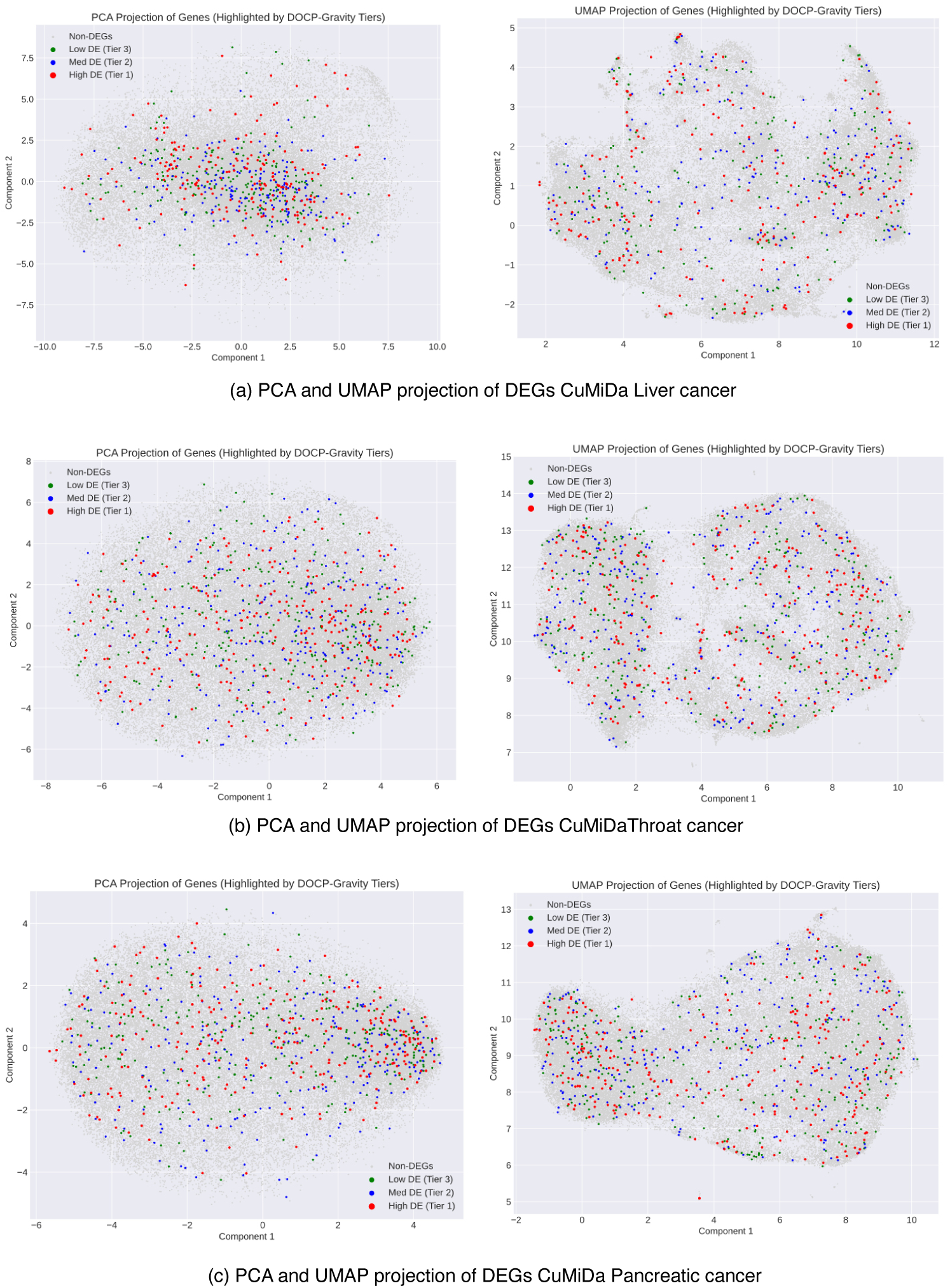
Visualisation of DEG distribution and UMAP projection of CuMiDa liver, throat and pancratic cancer datasets.

The PCA and UMAP based projections of the DEG feature space generated from the three CuMiDa cohorts, liver, throat and pancreas also indicate that the proposed DEG selection framework preserves discriminative expression profiles from microarray platforms as well. The distinguishable expression variance profile has been preserved accorss heterogeneous cancer cohorts while reduction of the influence from high-dimensional noise inherent to microarray data has been exhibited by the projections. In the CuMiDa cohort experimental pipeline, the selected DEGs produces a manifold clear boundary delination and reduced class overlapping. Particularly within the non-linear UMAP space. The preservation of coherent neighbourhood topology across both linear PCA and non-linear UMAp embedding suggests the selected DEGs may have retained biologically relevant expression relationships and underlying disease centric expression patterns, thereby supporting the feasibility for robust downstream tasks such as disease subtype detection and biomarker prediction. The PCA and UMAP representations of the CuMiDa microarray cohorts indicate that the proposed DEG selection framework preserves discriminative transcriptomic variance across heterogeneous expression profiles while reducing the influence of high-dimensional noise inherent to microarray platforms. In the Liver, Throat, and Pancreatic cancer datasets, the DEG-derived manifolds exhibit improved boundary delineation and reduced inter-class overlap between tumor and control samples, particularly within the non-linear UMAP space. Furthermore, the preservation of coherent neighborhood topology across both linear and non-linear embeddings suggests that the selected DEGs retain biologically relevant expression relationships and underlying disease-associated transcriptional patterns, thereby supporting their suitability for robust downstream classification and biomarker discovery tasks.

### Comparative Analysis

This sections presents comparative analysis of DEG selection performance through classification accuracy using three existing methods for DEG prediction from transcriptomic/microarray datasets. Two parametric methods, DESeq2 and Voom-Limma were used, and one non-parametric method NOISeq was used for benchmark comparison. Table 7 demonstrates the comparative analysis for the TCGA cohorts, while table 8 showcases the same for CuMiDa cohorts.

**Table 7:** Accuracy comparison of SVM and XGBoost models using DESeq2, Voom-Limma, NOISeq, and the proposed DEG-based method on LUAD, HNSC, and BRCA datasets.

| Model | Accuracy Metric | LUAD | HNSC | BRCA |
| --- | --- | --- | --- | --- |
| SVM | Accuracy (DESeq2) | $0.9652 \pm 0.0073$ | $0.9639 \pm 0.00014$ | $0.9878 \pm 0.00003$ |
| | Accuracy (Voom-Limma) | $0.9545 \pm 0.0068$ | $0.9833 \pm 0.0007$ | 1.0000 |
| | Accuracy (NOISeq) | 1.0000 | $0.9478 \pm 0.0023$ | $0.9855 \pm 0.0004$ |
| | Accuracy (Proposed) | $0.9886 \pm 0.0002$ | $0.9758 \pm 0.0007$ | $0.9919 \pm 0.00002$ |
| XGBoost | Accuracy (DESeq2) | $0.8957 \pm 0.0006$ | $0.9495 \pm 0.0006$ | $0.9802 \pm 0.00005$ |
| | Accuracy (Voom-Limma) | $0.9091 \pm 0.0070$ | $0.8756 \pm 0.0063$ | $0.9511 \pm 0.0029$ |
| | Accuracy (NOISeq) | $0.8636 \pm 0.0007$ | $0.8967 \pm 0.0054$ | $0.9282 \pm 0.0026$ |
| | Accuracy (Proposed) | $0.9433 \pm 0.0027$ | $0.9333 \pm 0.0027$ | $0.9785 \pm 0.00007$ |

**Table 8:** Accuracy comparison of SVM and XGBoost models using DESeq2, Voom-Limma, NOISeq, and the proposed method on CuMiDa liver, throat, and pancreatic cancer datasets.

| Model | Accuracy Metric | CuMiDa Datasets |  |  |
| --- | --- | --- | --- | --- |
|  |  | Liver | Throat | Pancreatic |
| SVM | Accuracy (DESeq2) | $0.9594 \pm 0.0009$ | $0.8724 \pm 0.0043$ | $0.8724 \pm 0.0043$ |
| | Accuracy (Voom-Limma) | $0.9558 \pm 0.0009$ | $0.7143 \pm 0.0074$ | $0.8127 \pm 0.0129$ |
|  | Accuracy (NOISeq) | NIL | NIL | NIL |
| | Accuracy (Proposed) | $0.9040 \pm 0.0011$ | $0.7971 \pm 0.0118$ | $0.7055 \pm 0.0001$ |
| XGBoost | Accuracy (DESeq2) | $0.8273 \pm 0.0221$ | $0.7105 \pm 0.0091$ | $0.7105 \pm 0.0091$ |
| | Accuracy (Voom-Limma) | $0.7206 \pm 0.0135$ | $0.7105 \pm 0.0091$ | $0.7182 \pm 0.1830$ |
|  | Accuracy (NOISeq) | NIL | NIL | NIL |
| | Accuracy (Proposed) | $0.8933 \pm 0.0021$ | $0.8357 \pm 0.0042$ | $0.8455 \pm 0.0170$ |

The comparative evaluation against existing benchmark methods clearly demonstrates that the proposed framework achives highly competitive and in several cases, supereor classification performance relative to established differential expression identification methods including DESeq2, Voom-Limma and NOISeq. On the TCGA cohorts, the proposed method consistently exhibited strong predictive accuracy across LUAD, HNSC and BRCA datasets for both SVM and XGBoost classifiers, while maintaining lower variance across repeated evaluations. Notably the proposed method have outperformed DESeq2 and Voom-Limma in LUAD and BRCA using SVM achieving accuracies of 98.86% and 99.19% respectively. This indicates improved preservation of discriminative transcriptomics signatures. Although NOISeq achieved perfect accuracy for LUAD with SVM, its performance was inconsistent across other cohorts and classifiers and may well have overfitted. In the CuMiDa microarray datasets the evidence of the robustness of the proposed framework is even more evident. While NOISeq has completely failed to identify any suitable DEGs which can be statistically significant across all microarray cohorts, indicating probable data quality issues or normalization failures or sample insufficiency, the proposed method consistently identified significant informative subsets that enables stable and robust downstream classification. Furthermore the proposed method has substantially outperformed DESeq2 and Voom-Limma for XGBoost across all three cohorts, liver, throat and pancreatic datasets achieving accuracies higher than the ones achieved by the feature space produced by the existing benchmark methods. These findings suggest that the proposed framework exhibits improved cross-platform generalizability and an enhanced resilience to heterogeneous transcriptiomic/expression noise compared to traditional statistical DEG extraction strategies.

### External Validation

This section presents and discusses the external validation results. The external validation was performed on four different instances. The first two instances uses the DEGs identified by the proposed framework from TCGA LUAD and BRCA datasets. These DEGs were used to classify the Gene Expression Omnibus Lung cancer (GSE 19804) and Breast cancer (GSE 70794) datasets obtained from the CuMiDa repository. The remaining two instances used the DEG feature space identified by the proposed framework from the Liver and Pancreatic cancer microarray cohorts from CuMiDa to classify the TCGA LIHC (Liver cancer) and TCGA PAAD (Pancreatic cancer) RNA sequence star count datasets. Table 9 presents the results respectively.

**Table 9:** External validation performance using TCGA-identified and GSE-identified DEG sets across independent cancer datasets.

| Dataset / Model | Original Accuracy | Accuracy on External DEGs | Original F1 | F1 on External DEGs |
| --- | --- | --- | --- | --- |
| **External validation of GSE data using TCGA-identified DEGs** |  |  |  |  |
| Lung GSE19804 (SVM-RBF) | $0.9560 \pm 0.0013$ | $0.9548 \pm 0.0016$ | $0.9562 \pm 0.0013$ | $0.9539 \pm 0.0016$ |
| Lung GSE19804 (XGBoost) | $0.9846 \pm 0.0008$ | $0.9843 \pm 0.0007$ | $0.9846 \pm 0.0008$ | $0.9846 \pm 0.0006$ |
| GSE70947 Breast Cancer (SVM-RBF) | $0.8147 \pm 0.0017$ | $0.8099 \pm 0.0015$ | $0.8112 \pm 0.0018$ | $0.8051 \pm 0.0018$ |
| GSE70947 Breast Cancer (XGBoost) | $0.7989 \pm 0.0015$ | $0.8721 \pm 0.0011$ | $0.7986 \pm 0.0015$ | $0.8716 \pm 0.0011$ |
| **External validation of TCGA data using GSE-identified DEGs** |  |  |  |  |
| TCGA LIHC (SVM-RBF) | $0.9732 \pm 0.0002$ | $0.9718 \pm 0.0001$ | $0.9848 \pm 0.0001$ | $0.9840 \pm 0.0001$ |
| TCGA LIHC (XGBoost) | $0.9742 \pm 0.0002$ | $0.9651 \pm 0.0003$ | $0.9854 \pm 0.0001$ | $0.9805 \pm 0.0001$ |
| TCGA PAAD (SVM-RBF) | $0.9818 \pm 0.0001$ | $0.9781 \pm 0.0001$ | $0.9728 \pm 0.0001$ | $0.9889 \pm 0.0001$ |
| TCGA PAAD (XGBoost) | $0.9818 \pm 0.0001$ | $0.9781 \pm 0.0001$ | $0.9728 \pm 0.0001$ | $0.9889 \pm 0.0001$ |

The results from the external validation clearly exhibits that the DEGs identified by the proposed method using a cohort dataset produced by sequencing technology can retain strong predictive capability across other platforms like microarray as well. This indicates that the proposed method can demonstrate strong modality agnostic performance in terms of classification tasks. The inference can be drawn that the proposed method generates high biological reproducibility and cross dataset generalizability without significant information loss. In the lung cancer dataset the TCGA-derived DEGs achieved nearly identical performance on external GSE19804 cohort with a marginal decrement of SVM and XGBoost accuracy. Suggesting that the selected genes can capture robust disease-associated expression signatures independent of sequencing platform variations. Similarly in the reciprocal validation setting, the microarray derived DEGs maintained strong performance on the TCGA LIHC and PAAD cohorts while experiencing only a minimal reduction in accuracy and F1 score compared to the original feature space.

Notably, the Breast cancer external validation produced improved performance for XG-Boost where accuracy and F1 score increased from 79.89% to 87.21% and 79.86% to 87.16% respectively when external DEGs from the TCGA BRCA cohort was used. This indicates that the transferred feature space may provide superior discriminative ability between cancer and control samples as well as cancer subtype samples as well. The external validation analysis demonstrates that the DEGs identified by the proposed framework retain strong predictive capability across independent transcriptomic cohorts, indicating high biological reproducibility and cross-dataset generalizability. In the Lung cancer datasets, TCGA-derived DEGs achieved nearly identical performance on the external GSE19804 cohort, with SVM-RBF accuracy decreasing marginally from 0.9560 to 0.9548 and XGBoost from 0.9846 to 0.9843, suggesting that the selected genes capture stable disease-associated expression signatures independent of sequencing platform variations. Similarly, in the reciprocal validation setting, GSE-derived DEGs maintained strong performance on TCGA LIHC and PAAD cohorts, where only minimal reductions in accuracy and F1-score were observed relative to the original feature space. Overall the the reliable consistency of classification performance across various independent and heterogeneous GEO and TCGA datasets confirm that the proposed DEG selection framework identifies statistically robust genes which are also a representative and informative subsets suitable for further downstream tasks such as biomarker prediction and disease prognosis analysis. The proposed DEG selection framework yields strong cross-platform stability, robustness along with reduced susceptibility to dataset-specific overfitting.

### Biological Enrichment and Analysis

Merely statistical superiority of the selected DEG feature space does not necessarily mean that the selection is inherently significant biologically as well. To determine the biological significance of knowledge representation by the selected subset Gene Set Enrichment Analysis (GSEA) and Kyoto Encyclopedia of Genes and Genomes (KEGG) pathway analysis was performed on the selected DEGs for all the cancer cohorts used in the experimental pipeline. The results of the GSEA and KEGG pathway analysis is provided in Figures S1-S6 of the supplementary information. The findings from the GSEA and KEGG pathway analysis shows that across LUAD, HNSC and BRCA the identified DEGs consistently recovered hallmark cancer biology centric information that focuses on DNA damage response, cell-cycle control, apoptosis, and proliferative signaling. In LUAD, the strongest signals involves p53-linked DNA repair, cell-cycle checkpoints, T-cell mediated immune regulation and angiogenesis-associated processes. HNSC exhibits a similar center, expressing prominent enrichment for G2/M transfusion, stem/progenitor cell proliferation, DNA repair and oncogenic signaling pathways such as Notch, Hedgehog, RAS and p53. BRCA genes dominate p53-mediated DNA damage response, apoptotic regulation, cell-cycle phase transition, and growth-factor/ ECM related pathways. The KEGG patterns reinforce the same pattern, but largely mapping to broad cancer and signaling pathways rather than tissue specific mechanisms. This suggests that the DEG selection mechanism captures shared oncogenic circuitry across tumor types. It identifies process governing genomic instability and proliferative escape. However the high p-values suggest that these enrichments should be interpreted as biologically coherent trends. Overall the method for TCGA cohorts is suitable for identification of compact gene sets comprising cancer-relevant genes that reflect tumorigenic programs across heterogeneous TCGA cohorts.

The same enrichment analysis when performed on CuMiDa datasets using the genes identified from them by using the proposed method, the results indicated coherent and consistent outcomes. Across the CuMiDa panels of Liver, Throat and Pancreatic cancer, the DEGs identified from them recovered core and robust cancer associated programs. In the pancreatic set, the prominent thematic concentration has centered around immune-cell proliferation, endothelial/angiogenic activation, p53-linked cellular cycle control and major oncogenic signaling axes such as the TGF-beta, Hippo, Notch, VEGF, ErbB and PI3K-Akt. The pattern significantly suggests that the selected genes are capable of capturing both rumor-growth signaling and microenvironmental remodeling. The Throat dataset exhibited stronger attention on DNA damage checkpoint control, apoptosis, G2/M regulation, and receptor-mediated signaling. Alongside the KEGG pathway enrichment for non-homologous end joining, proteoglycans in cancerous systems, complement/coagulation and circadian rhythm. These findings indicate that the proposed framework is sensitive to genes involved in genomic instability, cell cycle restraint, and extracellular signaling. For the Liver cohort the enrichment was dominated by DNA double-strand break repair, RNA capping/surveillance, spindle checkpoint control, cytoskeletal organization and T-cell mediated immunity. While the KEGG signals in ABC transporters, amino acid pathways, mRNA degradation, leukocyte migration and Th17 differentiation. Overall the CuMiDa results supports the claim that the method captures robust cancer-relevant signature across independent heterogeneous cohorts.

Collectively, the biological analyses across TCGA and CuMiDa datasets demonstrate that the proposed DEG selection method consistently captures conserved cancer-associated mechanisms, including cell-cycle dysregulation, DNA damage response, apoptotic signalling, immune modulation, and oncogenic pathway activation. Despite dataset heterogeneity, the enrichment profiles remained biologically coherent across tissue types, indicating the robustness and generalizability of the approach. These findings support the utility of the method for identifying compact yet functionally relevant gene signatures across diverse cancer cohorts. From a macro perspective, the biological analyses across two different sequencing platforms and cohort sets for heterogeneous cancer types, namely TCGA and CuMiDa (Gene Expression Omnibus) the results show promising outcomes. The proposed DEG selection mechanism consistently captured conserved cancer-associated mechanisms, which included cel-cycle dysregulation, DNA damage response, apoptotic signalling, immunity modulation and oncogenic pathway activation. Despite dataset heterogeneity and multimodal datatype from different sequencing platform, the enrichment profiles remained meaningful, robustly biologically relevant and expressed coherent cancer-specific knowledge across tissue types, indicating generalizability of the approach. These findings support the utility of using quantum mechanical axioms to develop computational methods to identify and extract meaningful information from biological datasets.

## Conclusions

This work presents a quantum-inspired composite-kernel framework for identifying differentially expressed genes from high-dimensional transcriptomic data. Instead of relying only on conventional distributional assumptions, the proposed method represents gene-expression states through normal and cancer-associated components, projects them into a Hilbert-space formulation, and uses the induced geometric structure to guide DEG selection. By combining curvature-based information with a gravitational-search-inspired optimization strategy, the framework selects a compact gene subset while explicitly controlling representational loss. The results show that the selected DEG feature spaces retain strong discriminative information across both RNA-seq and microarray datasets. In the TCGA cohorts, the proposed method generally improved classification performance compared with the original full transcriptomic feature space, particularly for SVM with RBF kernel, while XGBoost also showed clear gains in several cases. The CuMiDa experiments further indicate that the method is not restricted to one data-generation platform, although the magnitude of improvement remains classifier- and dataset-dependent. Comparative benchmarking against DESeq2, Voom-Limma, and NOISeq showed that the proposed method is competitive with established DEG-identification approaches and, in several cases, provides more stable down-stream classification performance, especially across heterogeneous microarray cohorts.

The external validation experiments are particularly important because they show that DEG sets identified from one platform can preserve predictive performance on independent datasets generated from another platform. This suggests that the proposed framework captures disease-associated expression signatures that are not merely dataset-specific artifacts. The biological enrichment analysis further supports this interpretation. Across TCGA and CuMiDa cancer cohorts, the selected genes were associated with biologically meaningful processes such as cell-cycle regulation, DNA-damage response, apoptosis, immune modulation, angiogenesis, and major oncogenic signaling pathways. These findings indicate that the selected DEG subsets are not only statistically useful for classification, but also biologically relevant for cancer-related downstream analysis.

Overall, the study demonstrates that a geometry-aware, quantum-inspired DEG selection strategy can provide a compact, informative, and biologically coherent representation of transcriptomic data. The framework shows promise for biomarker discovery, disease classification, and cross-platform transcriptomic analysis. At the same time, further validation on larger independent cohorts, additional cancer types, and single-cell datasets will be important to assess its broader robustness, biological specificity, and clinical translational potential.

## Acknowledgements

M.G. acknowledges the NVIDIA Center of Excellence hosted by Presidency University for supporting the research and allowing us to use the computational resources for the major experimentation of the proposed framework. A.P.S acknowledges Anusandhan National Research Foundation (ANRF) and S.N. Bose National Centre for Basic Sciences for providing postdoctoral fellowship to support this research. This work used the computing resources of ‘PARAM Shakti’ at IIT Kharagpur and ‘PARAM Rudra’ at S.N. Bose National Centre for Basic Sciences which are a part of the National Supercomputing Mission (NSM), implemented by C-DAC and supported by the Ministry of Electronics and Information Technology (MeitY) and Department of Science and Technology (DST), Government of India.

## Data and Code Availability

The code and the complete pipeline is available publicly at https://github.com/Apsphy/ Hilbert-Space-Projection-for-DEG.git and the datasets are available publicly from the Xena browser (https://xenabrowser.net/datapages/) and CuMiDa (https://sbcb.inf.ufrgs.br/cumida).

## Author Contributions

A.P.S worked on the mathematical formalism and development of the composite kernel. M.G worked on the loss function modeling. M.G developed significant parts of the code and the comparative benchmark execution, A.P.S developed some minor parts of the code and made changes during the experiments, and executed it equally by running various experiments on different clusters and supercomputers. M.G and A.P.S contributed equally to the preparation of the manuscript and significant discussions during the workflow experiment. The authors also declare no conflict of interest during any stage of preparation of this manuscript.

