## Supplementary Document for "A Curvature Guided Composite Kernel Framework for Differential Gene Selection in Cancer Transcriptomics"

### Supporting Information

### List of Tables

S1 Core Parameters Used in the proposed method-based feature selection framework . . . . S7
S2 Classification Experiment Parameters for the Experimental Pipeline . . . . . S8
S3 Differential Expression Analysis Parameters for DESeq2, Voom-Limma, and NOISeq Pipelines . . . . S9

### List of Figures

S1 Gene set enrichment analysis and KEGG pathway enrichment analysis for DEGs selected from TCGA LUAD . . . . S10
S2 Gene set enrichment analysis and KEGG pathway enrichment analysis for DEGs selected from TCGA HNSC . . . . S11
S3 Gene set enrichment analysis and KEGG pathway enrichment analysis for DEGs selected from TCGA BRCA . . . . S12
S4 Gene set enrichment analysis and KEGG pathway enrichment analysis for DEGs selected from Gene Expression Omnibus Liver Cancer . . . . S13
S5 Gene set enrichment analysis and KEGG pathway enrichment analysis for DEGs selected from Gene Expression Omnibus Throat Cancer . . . . S14
S6 Gene set enrichment analysis and KEGG pathway enrichment analysis for DEGs selected from Gene Expression Omnibus Pancreatic Cancer . . . . S15

### Appendix A: Pseudo-code of Algorithms

**Algorithm 1** PROJECTION KERNEL

**Require:** Expression matrix $X_{\text{cpu}} \in \mathbb{R}^{N_{\text{samples}} \times N_{\text{genes}}}$, label vector $labels_t$, batch size $batch_{\text{genes}}$
**Ensure:** Kernel vector $K_c \in \mathbb{R}^{N_{\text{genes}}}$

$(N_{\text{samples}}, N_{\text{genes}}) \leftarrow \text{shape}(X_{\text{cpu}})$
**if** $|labels_t| < N_{\text{samples}}$ **then**
    **raise** VALUEERROR
**else if** $|labels_t| > N_{\text{samples}}$ **then**
    truncate $labels_t$ to the first $N_{\text{samples}}$ entries
**end if**
$K_c \leftarrow \mathbf{0} \in \mathbb{R}^{N_{\text{genes}}}$
$normal_{\text{mask}} \leftarrow (labels_t = 0)$
$cancer_{\text{mask}} \leftarrow (labels_t = 1)$
**for** $start \leftarrow 0$ **to** $N_{\text{genes}} - 1$ **step** $batch_{\text{genes}}$ **do**
    $end \leftarrow \min(start + batch_{\text{genes}},\ N_{\text{genes}})$
    $batch_{\text{idx}} \leftarrow start : end$
    $X_{\text{batch}} \leftarrow \text{toDevice}(X_{\text{cpu}}[:, batch_{\text{idx}}])$
    $X_{\text{normal}} \leftarrow X_{\text{batch}}[normal_{\text{mask}}]$
    $X_{\text{cancer}} \leftarrow X_{\text{batch}}[cancer_{\text{mask}}]$
    $kc_{\text{normal}} \leftarrow \sum(X^2_{\text{normal}})$ ▷ column-wise
    $kc_{\text{cancer}} \leftarrow \sum(X^2_{\text{cancer}})$ ▷ column-wise
    $K_c[batch_{\text{idx}}] \leftarrow kc_{\text{normal}} + kc_{\text{cancer}}$
**end for**
**return** $K_c$

**Algorithm 2** Estimate Curvature

**Require:** Kernel vector $kernel_{\text{values}}$, neighborhood size $k_{\text{neighbors}}$
**Ensure:** Normalized curvature vector $curv_{\text{norm}}$

$kc \leftarrow \text{flatten}(kernel_{\text{values}})$ on CPU
$n \leftarrow |kc|$
$(kc_{\text{sorted}}, idx_{\text{sorted}}) \leftarrow \text{SortAscending}(kc)$
$window \leftarrow k_{\text{neighbors}}$
$curv \leftarrow \mathbf{0} \in \mathbb{R}^n$
**for** $pos \leftarrow 0$ **to** $n-1$ **do**
    $left \leftarrow \max(0,\ pos - window)$
    $right \leftarrow \min(n,\ pos + window + 1)$
    $neighbors \leftarrow kc_{\text{sorted}}[left : right]$
    $center \leftarrow kc_{\text{sorted}}[pos]$
    $dists \leftarrow |neighbors - center|$
    **if** $|dists| > 1$ **then**
        $local_{\text{var}} \leftarrow \text{Var}(dists)$
    **else**
        $local_{\text{var}} \leftarrow 0$
    **end if**
    $curv[pos] \leftarrow -local_{\text{var}}$
**end for**
$curv_{\text{unsorted}} \leftarrow \mathbf{0} \in \mathbb{R}^n$
$curv_{\text{unsorted}}[idx_{\text{sorted}}] \leftarrow curv$
$\mu \leftarrow \text{Mean}(curv_{\text{unsorted}})$
$\sigma \leftarrow \text{Std}(curv_{\text{unsorted}}) + 10^{-8}$
$curv_{\text{norm}} \leftarrow \dfrac{curv_{\text{unsorted}} - \mu}{\sigma}$
**return** $curv_{\text{norm}}$

**Algorithm 3** BUILD NEIGHBOR INDICES

**Require:** Kernel vector $kernel_{\text{values}}$, neighborhood size $k_{\text{neighbors}}$
**Ensure:** Neighbor index matrix $neighbors_{\text{unsorted}}$

$kc \leftarrow \text{flatten}(kernel_{\text{values}})$ on CPU
$n \leftarrow |kc|$
$(kc_{\text{sorted}}, idx_{\text{sorted}}) \leftarrow \text{SortAscending}(kc)$
$window \leftarrow \lfloor k_{\text{neighbors}}/2 \rfloor$
$neighbors \leftarrow \mathbf{0} \in \mathbb{Z}^{n \times k_{\text{neighbors}}}$
**for** $pos \leftarrow 0$ **to** $n-1$ **do**
    $left \leftarrow \max(0,\ pos - window)$
    $right \leftarrow \min(n,\ pos + window + 1)$
    $window_{\text{idx}} \leftarrow [left,\ left+1,\ \ldots,\ right-1]$
    **if** $|window_{\text{idx}}| < k_{\text{neighbors}}$ **then**
        $pad \leftarrow k_{\text{neighbors}} - |window_{\text{idx}}|$
        **if** $left = 0$ **then**
            $extra \leftarrow [right,\ right+1,\ \ldots,\ right+pad-1]$
        **else**
            $extra \leftarrow [\max(0,\ left-pad),\ \ldots,\ left-1]$
        **end if**
        $window_{\text{idx}} \leftarrow window_{\text{idx}} \,\|\, extra$
    **end if**
    truncate $window_{\text{idx}}$ to the first $k_{\text{neighbors}}$ entries
    **for** $j \leftarrow 0$ **to** $k_{\text{neighbors}}-1$ **do**
        $neighbors[pos,\ j] \leftarrow idx_{\text{sorted}}[window_{\text{idx}}[j]]$
    **end for**
**end for**
$neighbors_{\text{unsorted}} \leftarrow \mathbf{0} \in \mathbb{Z}^{n \times k_{\text{neighbors}}}$
$neighbors_{\text{unsorted}}[idx_{\text{sorted}}] \leftarrow neighbors$
**return** $neighbors_{\text{unsorted}}$

**Algorithm 4** GRAVITY OPTIMIZE

**Require:** Kernel vector $kernel_{\text{values}}$, curvature vector $curvature$, neighbor matrix $neighbors_{\text{idx}}$, iterations $n_{\text{iterations}}$, initial gravity constant $G_0$, decay parameter $\lambda$, event horizon scale $event_{\text{horizon\_scale}}$, computation device $device$, random seed $random_{\text{state}}$

**Ensure:** Selected gene indices $selected_{\text{idx}}$, optimal gene count $k_{\text{opt}}$, masses $M$, latent positions $X$

$kc \leftarrow \text{flatten}(kernel_{\text{values}})$ on $device$
$curv \leftarrow \text{flatten}(curvature)$ on $device$
$neighbors \leftarrow neighbors_{\text{idx}}$ on $device$
$n \leftarrow |kc|$
$curv_{\text{abs}} \leftarrow |curv|$
$M \leftarrow \dfrac{curv_{\text{abs}} - \min(curv_{\text{abs}})}{\max(curv_{\text{abs}}) - \min(curv_{\text{abs}}) + 10^{-8}}$
$M \leftarrow M + 0.05$
set random seed to $random_{\text{state}}$
$X \leftarrow 0.1 \cdot \text{Randn}(n, 3)$
$V \leftarrow \mathbf{0}$ with the same shape as $X$
**for** $t \leftarrow 0$ **to** $n_{\text{iterations}} - 1$ **do**
    $G_t \leftarrow G_0 \cdot e^{-\lambda t / n_{\text{iterations}}}$
    $X_i \leftarrow X$ expanded along a neighbor axis
    $X_j \leftarrow X[neighbors]$
    $direction \leftarrow X_j - X_i$
    $dist_{\text{sq}} \leftarrow \sum(direction^2) + 10^{-6}$
    $d_{kc} \leftarrow |kc[neighbors] - kc| + 10^{-2}$
    $m_i \leftarrow M$ expanded along a neighbor axis
    $m_j \leftarrow M[neighbors]$
    $force_{\text{mag}} \leftarrow G_t \cdot \dfrac{m_i \cdot m_j}{d_{kc} \cdot dist_{\text{sq}}}$
    clamp $force_{\text{mag}}$ above by 5.0
    $F \leftarrow \sum(force_{\text{mag}} \cdot direction)$ over neighbors
    $A \leftarrow F/M$ expanded along the coordinate axis
    $V \leftarrow 0.85V + 0.15A$
    $X \leftarrow X + V$
**end for**
$center_{\text{of\_mass}} \leftarrow \dfrac{\sum(X \cdot M)}{\sum M}$
$dists \leftarrow \|X - center_{\text{of\_mass}}\|$
$top_{\text{mass\_idx}} \leftarrow \text{TopK}(M,\ \max(10, \lfloor 0.01n \rfloor))$
$R_{\text{limit}} \leftarrow \text{Quantile}_{0.90}(dists[top_{\text{mass\_idx}}]) \cdot event_{\text{horizon\_scale}}$
$selected_{\text{mask}} \leftarrow (dists \leq R_{\text{limit}})$
$selected_{\text{idx}} \leftarrow \text{Where}(selected_{\text{mask}})$
$k_{\text{opt}} \leftarrow |selected_{\text{idx}}|$
**return** $selected_{\text{idx}}$, $k_{\text{opt}}$, $M$, $X$

### Appendix B: Parameter Description

Table S1: Core Parameters Used in the proposed method-based feature selection framework

| Parameter | Value | Role in the Method |
|---|---|---|
| KNN_CURVATURE | 200 | This parameter defines the neighbourhood size during curvature estimation. For each gene, the variance of distance is calculated within its local neighbourhood to estimate local geometric curvature. A larger value indicates smoother and more globally stable curvature estimation, whereas a smaller value increases local sensitivity. |
| N_GRAVITY_ITERS | 50 | This parameter defines the number of iterations used in the gravitational-search-based gene selection method. During each iteration, gene interactions based on simulated gravitational forces in latent space are calculated, leading to progressive refinement of the spatial organization of informative genes. Higher values allow more stable convergence of the optimization dynamics. |
| GRAVITY_K_NEIGHBORS | 70 | This parameter determines the number of nearest neighbouring genes participating in the gravitational interaction for each gene. It helps build the interaction graph between genes and controls the local interaction topology of the optimization process. Larger values increase interaction density and global cohesion, while smaller values emphasize localized structural relationships. |

Table S2: Classification Experiment Parameters for the Experimental Pipeline

| **Classifier** | **Parameter** | **Value** |
|---|---|---|
| **SVM** | Kernel | RBF |
| | $C$ (Regularization) | $10^{-2}$ |
| | Gamma | $10^{-3}$ |
| | Number of Iterations | 50 |
| | K-fold Cross Validation | 10 |
| **XGBoost** | Number of Estimators | 50 |
| | Learning Rate | 0.01 |
| | Booster | `gbtree` |
| | Max Depth | 5 |
| | Number of Iterations | 50 |
| | K-fold Cross Validation | 10 |

Table S3: Differential Expression Analysis Parameters for DESeq2, Voom-Limma, and NOISeq Pipelines

| Method | Parameter | Value / Role |
|---|---|---|
| **Voom-Limma** | Normalization Method | TMM normalization using `edgeR::calcNormFactors()` |
| | DEG Significance Threshold | Benjamini-Hochberg adjusted p-value $< 0.05$ |
| | Fold Change Threshold | $\lvert \log_2 \text{FC} \rvert > 1.0$ |
| | Statistical Model | Linear modelling using `lmFit()` with empirical Bayes moderation using `eBayes()` |
| | DEG Extraction Strategy | Pairwise contrasts for multi-class datasets followed by union of significant DEGs |
| | Transformation | Voom $\log_2$-CPM precision-weighted transformation |
| **NOISeq** | Probability Threshold ($q$) | 0.8 for DEG identification |
| | Normalization Method | TMM normalization using `norm = "tmm"` |
| | Number of Iterations ($r$) | 20 |
| | CPM Filtering Threshold | CPM $\geq 1$ in at least one group |
| | Pseudocount Parameter ($k$) | 0.5 |
| | Adjustment Parameter ($adj$) | 1.5 |
| | Random Seed | 42 |
| | DEG Extraction Strategy | Pairwise comparisons across classes followed by union of DEGs |
| **DESeq2** | Normalization Method | Median-of-ratios normalization |
| | DEG Significance Threshold | Benjamini-Hochberg adjusted p-value $< 0.05$ |
| | Fold Change Threshold | $\lvert \log_2 \text{FC} \rvert > 1.0$ |
| | Dispersion Estimation | Negative binomial dispersion modelling |
| | Statistical Test | Wald significance test |
| | DEG Extraction Strategy | Pairwise differential analysis with union of DEGs for multi-class datasets |
| | Random Seed | 42 |

### Appendix C: Gene Set Enrichment and KEGG Pathway Analysis Figures

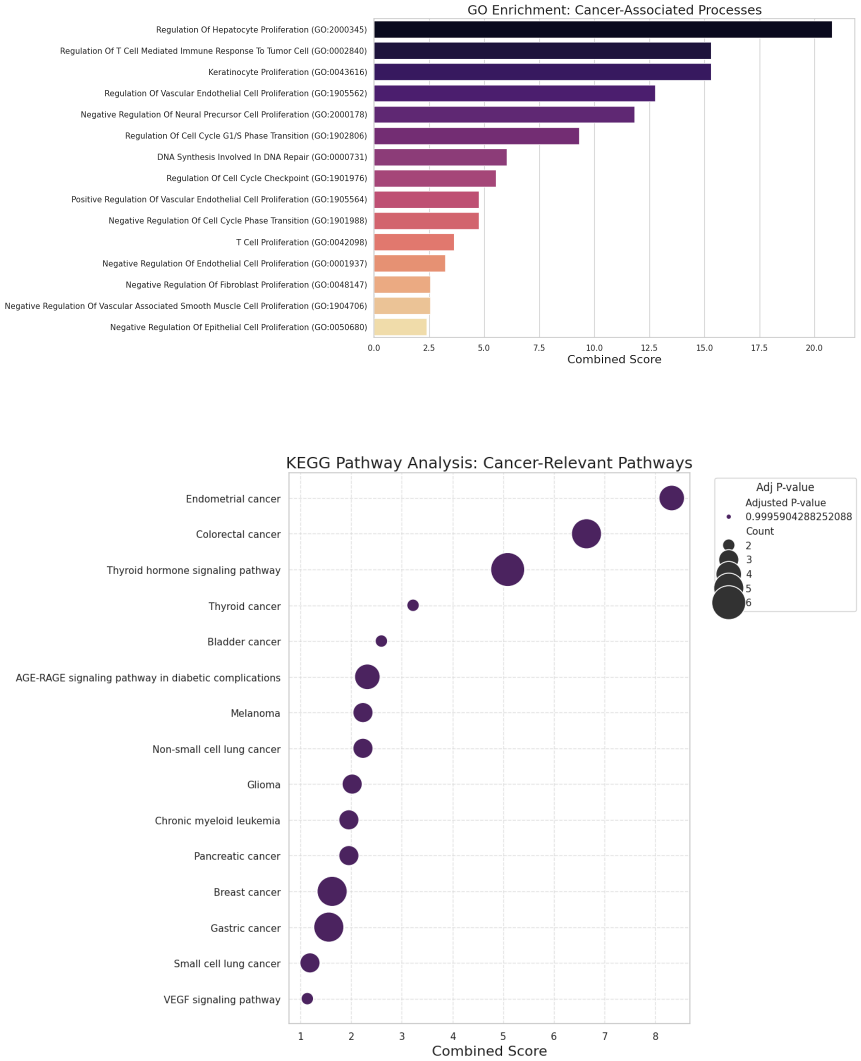


Figure S1: Gene set enrichment analysis and KEGG pathway enrichment analysis for DEGs selected from TCGA LUAD

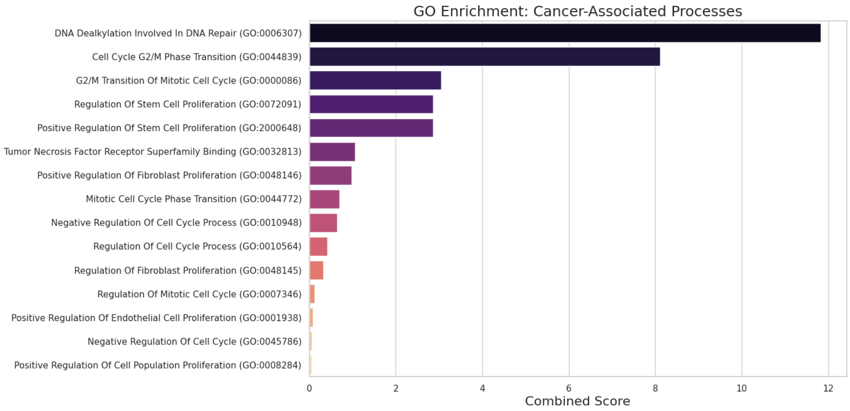


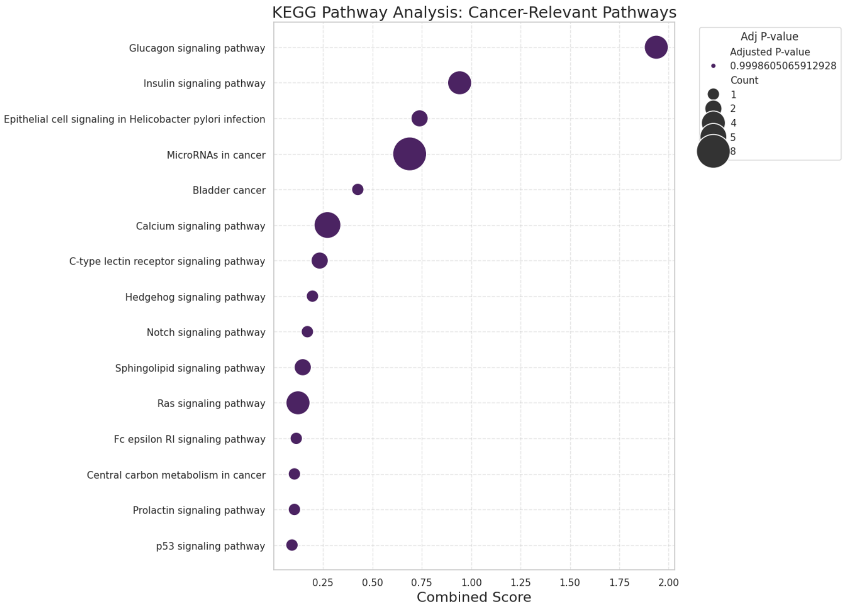


Figure S2: Gene set enrichment analysis and KEGG pathway enrichment analysis for DEGs selected from TCGA HNSC

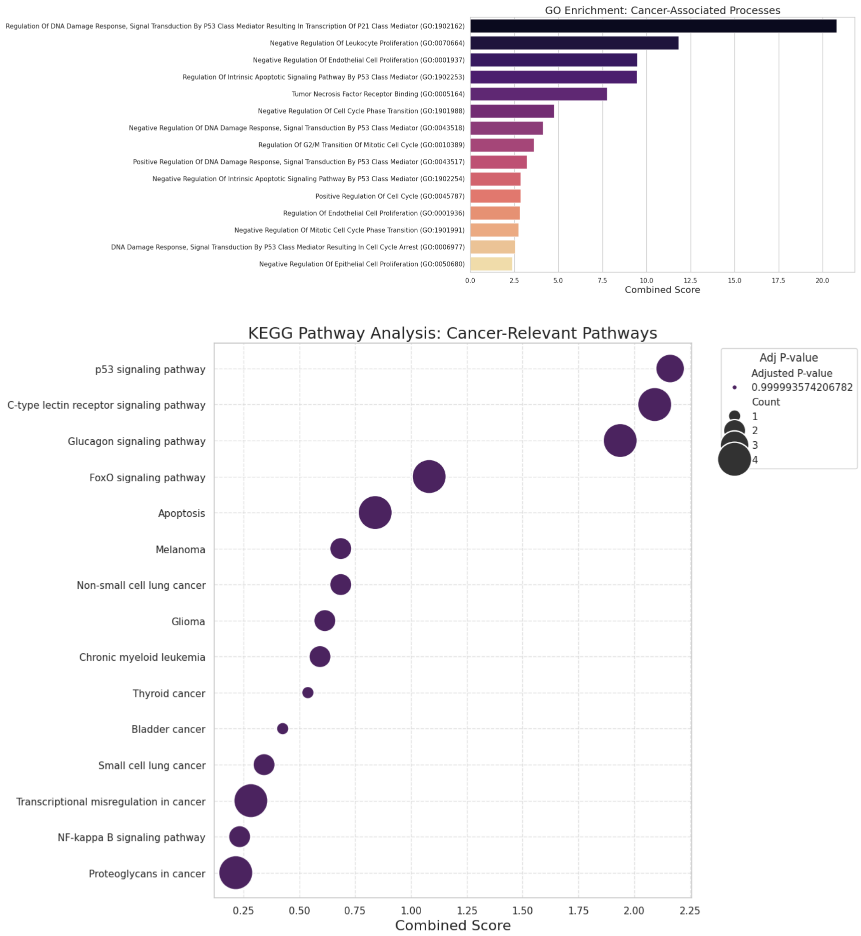


Figure S3: Gene set enrichment analysis and KEGG pathway enrichment analysis for DEGs selected from TCGA BRCA

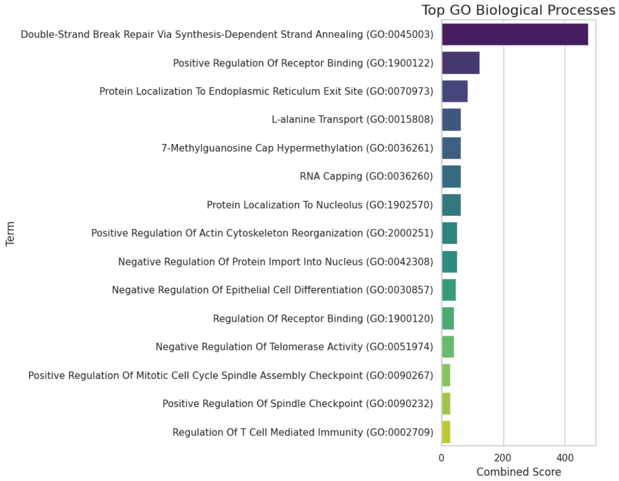


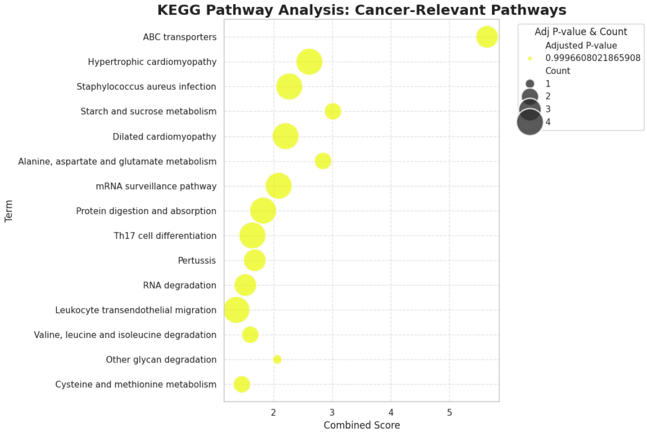


Figure S4: Gene set enrichment analysis and KEGG pathway enrichment analysis for DEGs selected from Gene Expression Omnibus Liver Cancer

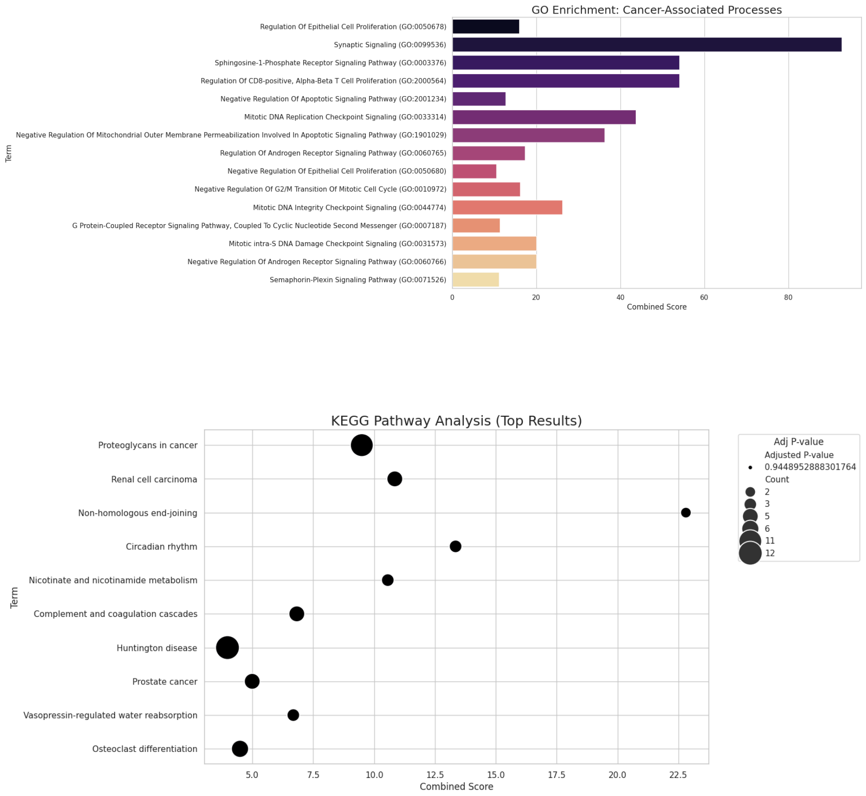


Figure S5: Gene set enrichment analysis and KEGG pathway enrichment analysis for DEGs selected from Gene Expression Omnibus Throat Cancer

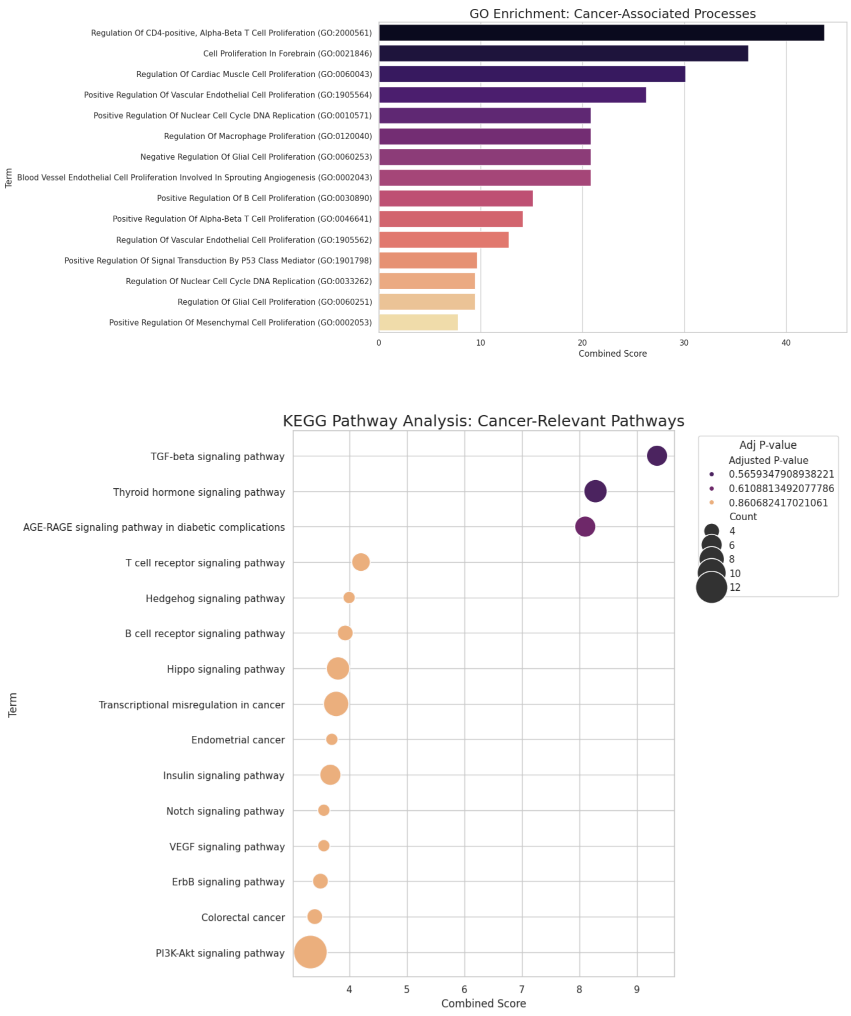


Figure S6: Gene set enrichment analysis and KEGG pathway enrichment analysis for DEGs selected from Gene Expression Omnibus Pancreatic Cancer
